# NORDIC Increases the Sensitivity and Preserves the Spatiotemporal Precision of fMRI Responses

**DOI:** 10.1101/2021.08.26.457833

**Authors:** Logan T. Dowdle, Luca Vizioli, Steen Moeller, Mehmet Akçakaya, Cheryl Olman, Geoffrey Ghose, Essa Yacoub, Kâmil Uğurbil

## Abstract

As the neuroimaging field moves towards detecting smaller effects at higher spatial resolutions, and faster sampling rates, there is increased attention given to the deleterious contribution of unstructured, thermal noise. Here, we critically evaluate the performance of a recently developed reconstruction method, termed NORDIC, for suppressing thermal noise using datasets acquired with various field strengths, voxel sizes, sampling rates, and task designs.

Following minimal preprocessing, statistical activation (t-values) of NORDIC processed data was compared to the results obtained with alternative denoising methods. Additionally, we examined the consistency of the estimates of task responses at the single-voxel, single run level, using a finite impulse response (FIR) model. To examine the potential impact on effective image resolution, the overall smoothness of the data processed with different methods was estimated. Finally, to determine if NORDIC alters or removes important temporal information, we employed an exhaustive leave-p-out cross validation approach, using FIR task responses to predict held out timeseries, quantified using R^2^.

After NORDIC, the t-values are increased, an improvement comparable to what could be achieved by 1.5 voxels smoothing, and task events are clearly visible and have less cross-run error. These advantages are achieved in the absence of large changes in estimates of spatial smoothness. Cross-validated R^2^s based on the FIR models show that NORDIC is not measurably distorting the temporal structure of the data and is the best predictor of non-denoised time courses. The results demonstrate that analyzing 1 run of data after NORDIC produces results equivalent to using 2 to 3 original runs and that NORDIC performs equally well across a diverse array of functional imaging protocols.

**Significance Statement:** For functional neuroimaging, the increasing availability of higher field strengths and ever higher spatiotemporal resolutions has led to concomitant increase in concerns about the deleterious effects of thermal noise. Historically this noise source was suppressed using methods that reduce spatial precision such as image blurring or averaging over a large number of trials or sessions, which necessitates large data collection efforts. Here, we critically evaluate the performance of a recently developed reconstruction method, termed NORDIC. Across datasets varying in field strength, voxel sizes, sampling rates, and task designs, NORDIC produces substantial gains in data quality. Both conventional t-statistics derived from general linear models and coefficients of determination for predicting unseen data are improved, while avoiding meaningful increases in typical estimates of image smoothness or substantial losses of temporal information.

## Introduction

The growing use of high (3 to 7 Tesla) and ultrahigh field (UHF, defined as ≥7 Tesla) magnetic fields for functional magnetic resonance imaging (fMRI) of brain activity has led to increases in the available signal-to noise-ratio (SNR) and subsequently corresponding interest in smaller voxel sizes and/or shorter repetition times. Early fMRI experiments tended to sample units of tissue that were on the order of 30 µL in voxel volume, a volume that potentially contains millions of neurons. In contrast, contemporary UHF high resolution fMRI studies, enabled by the significantly higher SNR and functional contrast-to-noise ratios (fCNR) available at UHF (Uğurbil, 2018, 2014), have attained resolutions that have ∼0.5 µL voxel volumes (e.g. ∼0.8 mm isotropic voxel dimensions). These developments, together with the early demonstration that neurovascular coupling has specificity at the level of mesoscopic scale organizations of the brain (Duong et al., 2001; Ugurbil, 2016), have led to a series of fMRI studies on cortical columns and layers (reviews (De Martino et al., 2018; Dumoulin et al., 2018; Finn et al., 2021; Lawrence et al., 2019; Norris and Polimeni, 2019; Polimeni and Uludağ, 2018; Weldon and Olman, 2021; Zaretskaya, 2021)). The use of high resolution functional imaging, largely initiated by the imaging of orientation domains together with ocular dominance columns for the first time in the human brain (Yacoub et al., 2008) and other fine scale organizations (e.g. (Huber et al., 2020; Stringer et al., 2011)) is growing more common.

The small voxel volumes in such high-resolution fMRI studies, however, have pushed the SNR of individual images and consequently the temporal SNR (tSNR) of the fMRI time series into a low SNR regime where the detectability of the functional responses become a major challenge. With this low SNR the thermal noise of the MR measurement begins to dominate the tSNR over signal fluctuations induced by physiological processes (often referred to as “physiological noise”) (Triantafyllou et al., 2011, 2005). Similarly, the use of highly accelerated fMRI approaches, introduced for rapid coverage of large volumes at high spatial resolution using UHF (Moeller et al., 2010; Uğurbil et al., 2013), has increasingly become the method of choice for data acquisition. These methods, popularized by the Human Connectome Project (Smith et al., 2013; Uğurbil et al., 2013), also push gradient echo (GE)-based fMRI data towards the thermal noise-dominated regime as repetition times and, consequently, flip angles and the signal magnitude detected in each image decreases (Smith et al., 2013; Uğurbil et al., 2013). While rapid sampling has led to a greater understanding of neural functioning and the BOLD response (Dowdle et al., 2021a; Polimeni and Lewis, 2021), it is not without consequences. The spatially non-uniform noise amplification introduced by parallel imaging reconstructions (i.e., the g-factor noise (Pruessmann et al., 1999)) further exacerbates the thermal noise penalty (Pruessmann et al., 1999; Todd et al., 2017). The resulting low SNR regime leads to difficulties in estimating the fine scale detail of the hemodynamic response, a critical goal given the variability of the hemodynamic response across large (Gonzalez-Castillo et al., 2012; Handwerker et al., 2004; Taylor et al., 2018) and small (Warren et al., 2014) regions of the brain. While there are potential statistical benefits for thermal noise dominance in meeting parametric assumptions in fMRI analyses (Wald and Polimeni, 2017), most researchers aim to remove it.

Unfortunately, the thermal noise associated with the MR measurement is not directly targeted by the various denoising approaches intended to suppress the contributions of structured, i.e. non-white, noise in an fMRI time series, emanating from physiological processes (e.g., (Bianciardi et al., 2009; Glover et al., 2000; Hu and Kim, 1994; Kay et al., 2013; Lund et al., 2006; Shmueli et al., 2007)), low-frequency signal drift, or motion. Spatial filtering (i.e. “smoothing”), on the other hand, does reduce the thermal noise contribution and hence is a commonly used approach to improve tSNR (Triantafyllou et al., 2006), and is a valuable approach when not fine detail is not desired (Wald and Polimeni, 2017). However, the addition of smoothing is often undesirable in high resolution fMRI as it results in substantial losses in spatial precision (Triantafyllou et al., 2006). Similarly, combining data from several different subjects reduces noise in general *via* group averaging; however, this option is not a desirable approach for high resolution studies because it inevitably incurs some degree of spatially non-uniform complex blurring nor is it valid for studies focused on single subject responses or inter-subject variability. Notably, in single subject statistical analysis approaches, such as multivoxel pattern analyses (MVPA) (Haxby et al., 2014) or encoding models (Kay et al., 2008; Naselaris et al., 2011; Vu et al., 2011), separate runs are typically leveraged in order to determine cross validated accuracy. In SNR-starved regimes in which neither smoothing nor group averaging are possible, these methods become difficult to implement, thereby limiting the range of possible scientific questions that can be addressed.

The growing focus on more spatially and temporally precise fMRI measurements, has recently led to the development of a denoising method, Noise Reduction with Distribution Corrected (NORDIC) PCA (Moeller et al., 2021; Vizioli et al., 2021). NORDIC suppresses Gaussian distributed noise associated with the MR detection process in repetitively acquired images, reducing thermal noise contributions throughout the image. The goal of NORDIC is to focus only on removing components of the timeseries which cannot be distinguished from Gaussian distributed noise, leaving the aforementioned non-white noise sources, such as physiological effects, signal drift or head motion as well as signals of interest, intact.

Prior work (Vizioli et al., 2021) with NORDIC primarily focused on submillimeter 7T fMRI data with an eye towards examining the functional point spread on the cortical surface of the primary visual cortex in response to a block design. The findings on such data were encouraging, suggesting no loss in functional precision or signal magnitude. However, it remained unclear if those findings would generalize to other fMRI acquisition paradigms and sequences. In this work, we further evaluate NORDIC’s utility as a denoising method for fMRI and compare it to a number of other noise suppression approaches using a variety of different datasets, which vary in field of view (up to whole brain), voxel size (0.8 to 2mm), repetition time (0.35 to 2.1s), field strength (3 and 7 Tesla), and use both block and event related task designs. The results obtained on 8 data sets (obtained from 3 subjects) provide a more detailed analysis of the NORDIC method and its generalizability. We find that NORDIC consistently leads to substantial gains in fMRI under a conventional generalized least-squares (GLSQ) framework and produces better single-run, single-voxel hemodynamic response function (HRF) estimates. Critically these effects are achieved with negligible increases of estimates of image smoothness. Collectively our findings suggest that these benefits are obtained while minimally affecting the intrinsic information present in the fMRI signal.

## Methods

### Stimuli and Datasets

For this manuscript, a total of 8 datasets (DS1 – 8) obtained from 3 subjects were considered (1 subject scanned 6 times, 2 others scanned once each). Two datasets (DS1,2) were examined under a different analytical framework in prior work, while portions of 4 others (DS3,4,6,7) were used for visualization purposes in supplemental material (Vizioli et al., 2021), but otherwise not analyzed. These datasets were chosen for their variability across multiple dimensions including field strength (3 or 7 T), sequence parameters (e.g., varying TR or voxel size), type of experimental design (block vs event) and field of view. For all stimuli, participants viewed the images though a mirror attached to the head coil. Datasets 1, 2, 5, 6, and 7 (DS1, DS2, DS5, DS6, and DS7) are block designs which used a flashing checkerboard (8Hz) positioned either in a center position (‘target’) or in a surround with the center cut out (“surround”), centered on a gray background. The center stimulus subtended approximately 6.5 degrees of visual angle, as did the width of the surround border. Stimuli were presented in a standard 12 s on 12 s off block design paradigm. At 7T, stimuli were presented on a Cambridge Research Systems BOLDscreen 32 LCD monitor positioned at the head of the scanner bed (resolution 1920 x 1080 at 120 Hz), whereas at 3T the stimuli were presented using a NEC NP4000 projector, using a projection screen placed at the end of the bore of the MR scanner (resolution 1024 x 768 at 60 Hz).

Dataset 3 (DS3) used an event related design in which full-color, intact and phase (of the image content) scrambled faces were presented. Stimuli were centered on a gray background. Stimuli were on screen for 2 seconds and separated by at least a 2 s interstimulus interval (ISI). For all runs, blank trials (2 per run) were included to jitter the stimulus presentation.

For Dataset 4 (DS4), we used an event related design with grayscale images of faces (20 male, 20 female) presenting neutral expressions. We manipulated the phase coherence of the image of each face from to produce 5 visual conditions, producing in 200 unique stimuli (5 visual conditions x 20 identities x 2 genders), as in previous work (Dowdle et al., 2021b). Stimuli approximately subtended 9° of visual angle. Stimulus presentation began and ended with a 12 s fixation period and had a duration of approximately 3 mins and 22 s. Within each run, we showed 40 images, each presented for 2 s, with a 2 s ISI as well as 10% blank trials (i.e., 4 s of fixation) randomly interspersed amongst the 40 images, effectively jittering the ISI.

For Dataset 8 (DS8), we used a modified rotating wedge retinotopy paradigm. The frequency of ring sweeps was approximately 0.05Hz. The subject maintained fixation on a central point throughout the task. The wedge extended from the central fixation point to the edge of the screen, with a width of 20 degrees.

For DS1-7, stimulus presentation was controlled using Psychophysics Toolbox (3.0.15)-based scripts on a Mac Pro Computer. For DS8, the stimulus was controlled using custom, inhouse developed software.

### MRI Acquisition

All functional MRI data were collected with either a 7T Siemens Magnetom System with a single channel transmit and 32-channel receive NOVA head coil or a 3T Siemens Magnetom Prisma^fit^ system using the Siemens 32-channel head coil. All functional images were obtained using T2*-weighted, simultaneous multislice (SMS)/multiband(MB) gradient echo, Echo Planar (GE-EPI) (Moeller et al., 2010) as developed and implemented in the Human Connectome Project (Uğurbil et al., 2013).

#### 7T fMRI Data (DS1 to DS5, and DS8)

For DS1 and DS2 imaging was restricted to the posterior occipital lobe, capturing 42 slices using a right to left phase encoding direction. DS3 images captured 42 slices of the occipital pole and ventral temporal lobe using an anterior to posterior phase encoding direction. For DS4 and DS5, whole brain images were collected, with 85 slices using an anterior to posterior phase encoding direction, with parameters matched to the HCP 7T Protocol. DS8 images captured only 32 slices of the occipital pole using a left to right phase encoding direction.

#### 3T fMRI Data (DS6 and DS7)

DS6 was acquired with a higher resolution than typically used at 3T studies (1.2 mm isotropic), capturing most of the brain with 100 slices, excluding the cerebellum, with anterior-to-posterior phase encoding.

DS7 was a whole brain study, 72 slices were acquired using an HCP-like 3T protocol. (See Table 1 for full details).

**Table 1.**
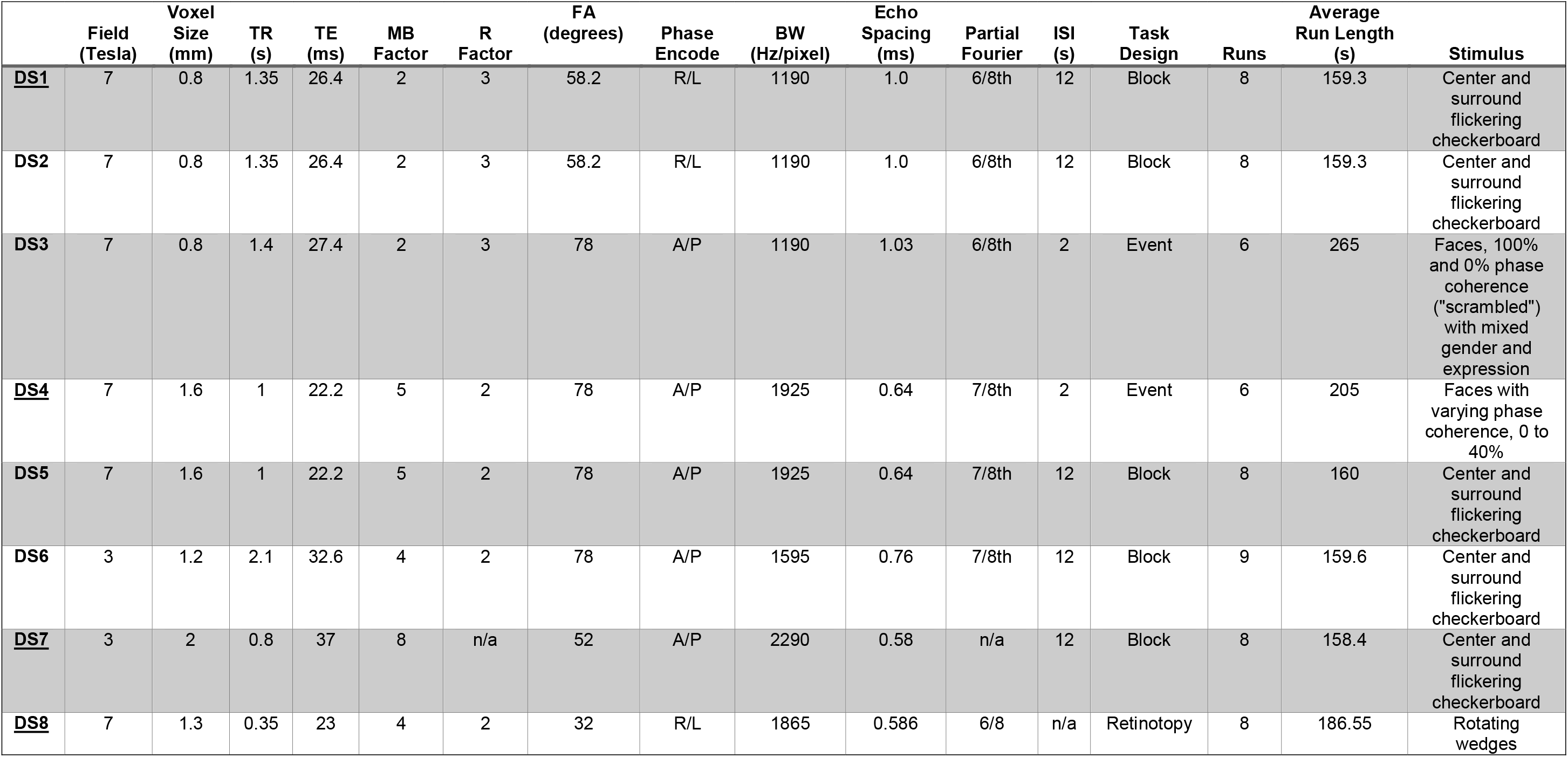
Dataset Acquisition and Task Details. Parameters are shown for all 8 datasets considered in the present work. TR: repetition time, TE: echo time, MB Factor: Multiband acceleration factor, R Factor: GRAPPA acceleration factor, FA: flip angle, BW: Bandwidth, ISI: Interstimulus interval, s: seconds.

#### Anatomical Imaging

T1-weighted anatomical images were obtained for DS1-DS7 using an MPRAGE (Mugler and Brookeman, 1991) sequence (192 slices; TR=1900 ms; FOV: 256 x 256 mm; flip angle= 9°; TE= 2.52 ms; spatial resolution 0.8 mm isotropic voxels) which were collected with a 3T Siemens Magnetom Prisma^fit^ system. The anatomical images for DS8 were acquired using a MP2RAGE (Marques et al., 2010) sequence (192 slices; TR=4300 ms; FOV: 240 x 225 mm; flip angle= 4°; TE= 2.27 ms; spatial resolution 0.8 mm isotropic voxels) collected with a 7T Siemens Magnetom scanner. Anatomical images were used only to visualize data and define regions of interest (ROIs, see Region of Interest Creation, below).

#### Initial MR Image Preprocessing

Two separate reconstruction methods were used in all subsequent analyses described below. Following data acquisition, the k-space data files for each receive channel produced by the SIEMENS system were saved. These were reconstructed *offline* (as opposed to using the scanners inbuilt reconstruction algorithms) using standard techniques (noise-decorrelation between channels, zero-filling for partial Fourier, split slice-GRAPPA for joint SMS and GRAPPA reconstruction (7×7 kernel), SENSE1 for multichannel combination with ESPIRIT calculated sensitivity profiles, and g-factor calculated from the SMS kernels and sensitivity profiles) implemented in-house to produce magnitude images with minimal processing, similar to the typical DICOM images produced by the default Siemens reconstruction. These minimally processed data are referred to as the “Standard” reconstruction, to emphasize the fact that this is a standard or typical reconstruction of the magnitude images.

The second image reconstruction, which is the primary consideration of this manuscript, is derived from the same raw k-space files and reconstruction steps, however, we applied additional denoising steps that aim to suppress thermal noise with NORDIC (Moeller et al., 2021; Vizioli et al., 2021). In brief, this method uses a patch based, PCA approach to identify and discard components of the data that are indistinguishable from zero-mean, normally distributed (i.e., thermal) noise, using the magnitude and complex portions of the MRI signal as input. NORDIC share similarities with existing low-rank methods (Candès et al., 2013; Haldar and Liang, 2011; Meyer, et al., 2020; Thomas et al., 2002; Veraart et al., 2016) but differs in key aspects. For example, NORDIC determines a noise threshold using noise estimates from the data itself and furthermore does so after correcting for spatial variability in noise (“g-factor”). In addition, NORDIC performs phase normalization, and is able to use larger patch sizes than comparable methods (Moeller et al., 2021; Vizioli et al., 2021). We used the default settings of NORDIC on all datasets, which maintains a minimum 11:1 ratio between spatial and temporal dimensions. In the datasets considered here this resulted in patches of size 11^3^ voxels for DS1,2; 12^3^ for DS3,5; 14^3^ for DS4, 7; 10^3^ for DS6 and 20^3^ for DS8. We refer to these data as “NORDIC” throughout the remainder of the manuscript. The use of an offline reconstruction for these data, rather than the typical scanner reconstruction assured that, other than the denoising step, all other reconstruction steps were identical.

In addition to the Standard and NORDIC reconstructions, we also considered a third approach referred to as dwidenoise (Cordero-Grande et al., 2019; Veraart et al., 2016) as provided with version 3.0.0 of MRtrix3 (Tournier et al., 2019), which was applied to the Standard magnitude images. In brief, dwidenoise, like NORDIC, also aims to suppress normally distributed noise using a patched-based denoising approach. Noise components for each patch are estimated on the basis of Marčenko-Pastur principal component analysis (MPPCA) which attempts to account for spatial variability in the noise. The recommended and validated default settings were used, with the size of the patch depending primarily on timeseries length. For DS1, DS2 and DS6, this led to a 5^3^ voxel patch size, whereas Datasets DS3, DS4, DS5 and DS7 had a 7^3^ patch size. We chose dwidenoise as a comparison as it is in active use and development, with a publicly available implementation. The “Standard” data was considered the reference point for further analyses.

Prior to any additional processing, we examined the noise removed by NORDIC and dwidenoise. Specifically, the noise removed by NORDIC was calculated by taking the mean of the magnitude of the complex difference between the Standard and NORDIC data. The noise removed by dwidenoise residuals was calculated as the absolute value of the mean difference between the Standard and dwidenoise data. Maps of the g-factor were derived from the k-space data files.

### Processing

All subsequent fMRI processing was performed using AFNI (Cox, 1996) and was identical for preparations of the data. For all datasets and denoising methods, we used conventional processing steps, with settings chosen to minimize any loss of precision. First, slice timing was corrected by using Fourier interpolation, with the first slice as the reference timepoint. Motion correction was then performed using the first volume of the first run from the Standard reconstruction as the registration target, with the ‘Fourier’ estimation and interpolation option chosen. Using the same target for motion correction across all reconstruction methods allows for subsequent voxel to voxel comparisons between the different methods. Regardless of whether we used standard or denoised data were used as input, the estimated motion parameters are highly similar with an average Pearson’s correlation coefficient > 0.99 (See Supplemental Table 2 for all values).

In order to compare the signal characteristics of NORDIC to more typical approaches that aim to reduce thermal noise, we created three additional comparator datasets from the standard data. These are 1) data smoothed with a FWHM gaussian kernel equivalent in size to one voxel (hereafter labeled “+1 voxel FWHM”), 2) data smoothed with a FWHM gaussian kernel equivalent in size to 1.5 voxels (“+1.5 voxel FWHM”), and 3) data temporally smoothed (“+temporal smoothing”) using a sliding window average approach with window sized between 9 and 10.5 s.

Data were then scaled voxel-wise to have a temporal mean intensity of 100 per run, which eases percent signal change calculations. In order to evaluate the fMRI statistical performance of each data set we considered two general linear modeling frameworks.

### Task Event Modeling

#### GLM One (Conventional Approach)

The scaled data were passed through a generalized least squares (GLSQ) regression model using a conventional hemodynamic response. Here we specifically used the double gamma hemodynamic response estimate provided with AFNI, “SPMG1”, with approximate peaks at 6 seconds (positive peak) and 17 seconds (negative undershoot). This was convolved with the stimulus time courses associated with each event type to produce the predictive model. GLSQ regression produced betas (i.e., parameter estimates/activation amplitudes) and t-statistics from the model fit for each event on a per run basis for each of the 9 (DS6), 8 (DS1, DS2, DS5, DS7), or 6 (DS3, DS4) runs. This was performed using AFNI’s *3dREMLfit*, which calculates an autoregressive moving average (ARMA(1,1)) model to estimate the temporal autocorrelation on a voxel by voxel basis, thereby improving the accuracy of t-statistic estimates (Olszowy et al., 2019).

#### GLM Two: Finite Impulse Response (FIR) Model

To investigate the temporal information present in each functional acquisition, we also used a finite impulse response (FIR) model using AFNI’s *3dDeconvolve* function with TENT estimators. These are ‘tent’ or ‘hat’ functions which are identical to delta functions when the stimuli rounded to each TR. The window for which these estimates were created varied between datasets, ranging from 15 to 29.4 seconds out from stimulus onset, but was identical between processing methods. This approach uses the repetition of identical stimuli within a run to estimate the voxel-by-voxel response to each stimulus class in a flexible manner, with no *a priori* assumptions regarding its specific shape.

The events for DS3 were separated by 1-second steps. Thus, the data for the FIR model was simultaneously slice time corrected and up-sampled to a 1-second sampling rate using in-house code prior to processing in AFNI. This step was performed in an identical manner for the Standard, NORDIC, and dwidenoise data, and was performed only for the FIR model. Though we show time courses for the HRF estimates for only the “Target” condition, the full model was used for cross validated prediction accuracy introduced below (see Temporal Precision).

### Region of Interest (ROI) Creation

#### Anatomical ROIs

Anatomical images for each dataset were segmented into different tissues and skull-stripped using the Segment tool from SPM12 (Ashburner and Friston, 2005). These skull-stripped anatomical images were aligned (rigid-body) to the mean of the first run of the standard data, after it had undergone motion correction, using a local Pearson correlation estimator (Saad et al., 2009) (*3dAllineate)*. We then applied the rigid-body transformations to the first three tissue class images, corresponding to gray matter, white matter and a mixture of CSF and vasculature (hereafter just CSF) such that they overlapped with the functional imaging data. The aligned tissue probability maps were then converted to binary masks, with a threshold of 0.95 probability for each class and then resampled to match each dataset’s EPI grid. These ROIs were then further restricted to voxels that contained sufficient functional image signal using a binary mask. This binary mask, automatically generated during processing, is a contiguous volume produced by an interactively clipping process which excludes the background and very low intensity values (*3dAutomask*) in the functional image. This mask will be subsequently described as the “EPI mask”. This masking was performed to minimize the amount of each anatomical ROI that includes voxels outside of the acquired field of view or overlapped with areas of near-complete signal dropout. These steps produced grey matter, white matter, and CSF regions of interest (ROIs) which are aligned to each unique functional dataset. Note that no distortion correction was applied to the functional data to minimize any additional blurring.

#### Functional ROIs

Multiple ROIs were created using all runs of the Standard data. We used the t-statistics derived from the GLSQ model’s fit corresponding to the contrasts of interest from each dataset. For all datasets (except DS4, DS8) we created a “Target ROI”, which was created by combining the multiple clusters with more than 10 contiguous activated (defined as contact via faces, edges, or corners) voxels using a voxel threshold of p<0.001 for contrast of the target stimulus (center or faces) vs the alternative stimulus (surround or scrambled faces).

To arrive at the minimum cluster size threshold of 10 voxels, we estimated the cluster size (number of voxels) required to obtain a cluster family wise error rate of p<0.05 using a Monte Carlo method as implemented in *3dClustSim*. This AFNI tool uses the smoothness estimates of the residuals to simulate 10,000 smoothness matched, noise-only datasets. This creates a create a null distribution of cluster sizes, from which a cluster size threshold can be obtained. This was done only for the Standard data and these cluster thresholds were only used to control false positives in the ROIs and not considered further. For all datasets and contrasts, 10 contiguous voxels at this threshold produced clusters that exceed the typical p_FWE_<0.05 threshold.

For DS4, which used face stimuli with variable phase coherence, the ROIs were generated from a separate localizer analysis of Standard data, using Faces greater than Scrambled Faces contrast. No functional ROIs were created for DS8, as it was only used for the frequency spectrum analysis (see Temporal Precision Evaluation).

In a similar way, a “Non-Target” ROI was also created by selecting all clusters greater than 10 contiguous voxels at a voxel-wise threshold of p<0.001 and positive signal associated with the alternative condition, that is, surround or scrambled faces, only. The use of these two different contrasts produced a complementary selection of voxels.

Collectively we produced 2 functional ROIs and 3 anatomically derived ROIs, per dataset. These were then used to summarize the distribution of the values from other voxel wise measures (e.g., t-statistics).

### Spatial Precision Evaluation

#### Global Smoothness

Spatial precision was estimated using smoothness estimates produced from each dataset within the previously described EPI mask. These smoothness estimates, produced by *3dFWHMx*, were conducted for three stages in processing and analysis. This measure is based on estimating the spatial autocorrelation function in each of the 3 voxel dimensions within a mask and reporting the average for the image volume. Any spatial smoothing introduced by post-acquisition data manipulations shows up as an increase in the estimate (in units: mm FWHM) relative to the Standard data. As this captures the average smoothness of the entire image volume, we refer to this as ‘global smoothness’. Specifically, we calculated global smoothness on each dataset prior to any processing, after processing and on the residuals from the conventional GLM. For the first two stages, the data were detrended (including removal of the mean) to remove temporal drifts and variation in voxel intensities related to anatomy. Significance was evaluated using paired t-tests between the 3 primary types of processing (Standard, NORDIC, dwidenoise).

#### Local Smoothness

In addition, we performed an analysis using the AFNI function *3dLocalACF*, which estimates a voxel-wise spatial autocorrelation function, using a spherical local neighborhood with a radius of 10 voxels. This tool examines each voxel within a brain mask and correlates its timeseries with that of its neighbors. The gaussian + mono-exponential autocorrelation function is then fit to the resulting map of Pearson’s correlations, to provide a voxel-by-voxel estimate of smoothness. We refer to this as local smoothness since the parameter is estimated on a voxel wise basis and is expected to highlight regional variations in image smoothness. As this method is highly sensitive to trends within the data, this local smoothness was estimated only on the residuals of the conventional GLM. This was done independently per run, and then averaged. Values, in mm FWHM, were then summarized in each of our three tissue masks: gray matter, white matter and CSF (See Region of Interest Creation).

Naturally, the GLM residuals used in both the global and local smoothness calculations will retain some structure not attributable to thermal noise and not captured by the task and nuisance regressors of the GLM, but in working with real-world data, this is the best available approximation to structure-free data.

### Temporal Precision Evaluation

#### Fourier Spectrum Analysis

Using DS7 and DS8, which were sampled at 800 and 350ms respectively, we performed a fast Fourier transform across time on each independent run of the data, doing this for the Standard and NORDIC data. The FFT was performed on the scaled data, which is the final output of the magnitude data preprocessing pipelines. To show the frequency spectrums and their variability within each tissue class we took the mean of the absolute value across voxels within each tissue class (GM, WM, CSF), and then the mean and standard deviation across each independent run. We then examined the normalized frequency spectrum within each ROI up the Nyquist frequencies (DS7: 0.625Hz, DS8: 1.429Hz) to determine how NORDIC processing altered the frequency spectrum and if physiological frequency peaks remained in the data. For DS7, the CSF, gray matter and white matter ROIs contained 13314, 66664 and 42356 voxels respectively. For DS8, the CSF, gray matter and white matter ROIs contained 2367, 51797 and 19281 voxels respectively.

#### Cross validation

We used an exhaustive Leave-p-Out permutation approach to evaluate the accuracy of the estimated HRF time courses derived from the full FIR model (“GLM Two”, see above) and to determine if the NORDIC denoising method altered their temporal structure. Specifically, we varied the number of runs, P which ranged between the total number of runs-1 (N-1) and 1 to be used as a test set and trained with the remaining runs. Prior to being entered into the model, we projected out polynomials (up to order = run duration in minutes - 1) to remove low frequency drift and masked the data using the EPI data mask derived from the Standard data.

In order to limit our analysis to voxels that were plausibly task responsive, we determined the overall leave-one-out cross validated coefficient of determination R^2^ using all runs of each Standard dataset in a model with a conventional HRF. This generated one map of R^2^ across the whole brain for each Dataset. This map was used only to summarize the subsequent exhaustive FIR based cross-validation scheme.

In each fold, we used N-P runs to estimate the FIR model, constructing a series of betas for each stimulus, corresponding to the estimated BOLD response over time to each stimulus on a voxel-by voxel basis. These estimates were then multiplied by a design matrix for the held-out runs (P) to generate predicted timeseries for each voxel. We then determined how well the predicted timeseries matched the true timeseries using the coefficient of determination, R^2^. We first considered the P=N-1 case, in which one run was used to predict the timeseries of all remaining runs. The R^2^ (and standard error over permutations) of the FIR cross-validation scheme was calculated for voxels ranging from 5% of variance explained in the conventional model to the max R^2^ for that dataset.

For 1≤P≤N-2, we summarized error across permutations using a R^2^≥15% mask from the conventional model. This process was repeated to generate the following three comparisons: Standard predicting Standard, NORDIC predicting NORDIC, and, most importantly, NORDIC predicting Standard. This final comparison is significant, as it represents the NORDIC data predicting the non-denoised data, which has undergone minimal processing, and retains all of the signals of interest, albeit mixed with noise.

## Results

Three representative datasets (of 8) were selected to present in figures. Unless indicated otherwise, the summary metrics provided in the results refer to the mean across 7 datasets (DS1-7; DS8 was only considered for FFT analyses), normalized if necessary (i.e., due to different voxel sizes).

### Conventional General Linear Model (GLM) results

Figure 1 shows the distribution of t-values from the conventional GLM analysis on all runs combined, using a canonical HRF. Processing with NORDIC leads to an increase in the t-values, visible as a large rightward shift in their distribution relative to the data reconstructed with the Standard data. The t-values were extracted from an identical ROI, created based on the Standard data (See Methods). This increase in t-values is found within both the Target ROI (DS1,DS6: center > surround checkerboard; DS3: faces > scrambled) as well as the non-Target ROI (DS1, DS6: response to surround checkerboard; DS3: scrambled stimuli only). This effect is consistent across all 7 Datasets (Supplemental Figure S1); the mean of the one-sided t-statistic across datasets within the non-target ROI was 8.7±5.0 for NORDIC and 5.67±3.6 for the Standard Reconstruction. In the Target ROI these values were 9.9±4.5 for NORDIC and 6.16±3.6 for Standard.

**Figure 1.**
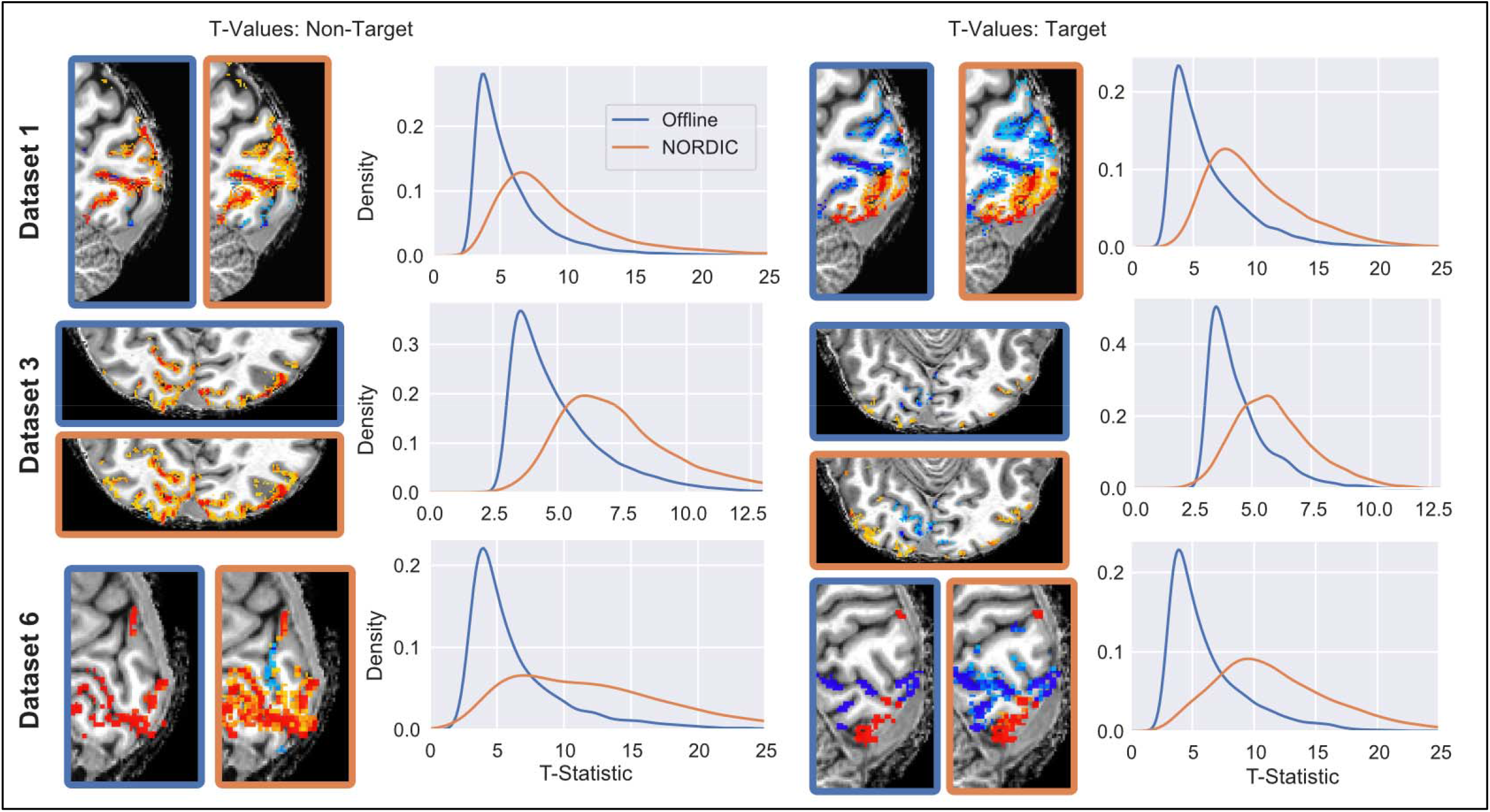
The distributions of t-Statistics from the model using all runs of data are shown. Distributions show the t-statistics of voxels from Non-Target (left) and Target (right) ROIs, defined as those that displayed significant positive stimulus-evoked changes relative to baseline (Non-Target) or in the contrast between Target and Non-Target conditions (Target) in the Standard data. Functional maps of the corresponding contrast are shown for visual reference at a t-value threshold of 3.3, corresponding to voxel-wise p<0.001 (uncorrected). NORDIC and Standard reconstructed functional maps are identified by the color of the border of the two images shown for each dataset (blue=Standard, orange=NORDIC). Across all datasets and both ROIs, the distribution of t-statistics for NORDIC was higher, with a longer tail.

### Comparison with Other Noise Reduction Methods

To evaluate the relative performance of NORDIC compared to other methods that seek to reduce normally distributed noise, we ran identical general linear models (GLMs) on data that were additionally processed with dwidenoise (Cordero-Grande et al., 2019; Veraart et al., 2016), spatially (1 and 1.5 voxels FWHM) or temporally (sliding window ∼10s) smoothed following preprocessing (See Methods). Here we only consider the Target ROI. T-Statistic distributions for DS1, 3, and 6 are shown in Figure 2. Results for all datasets are given in Supplementary Figure S2.

**Figure 2.**
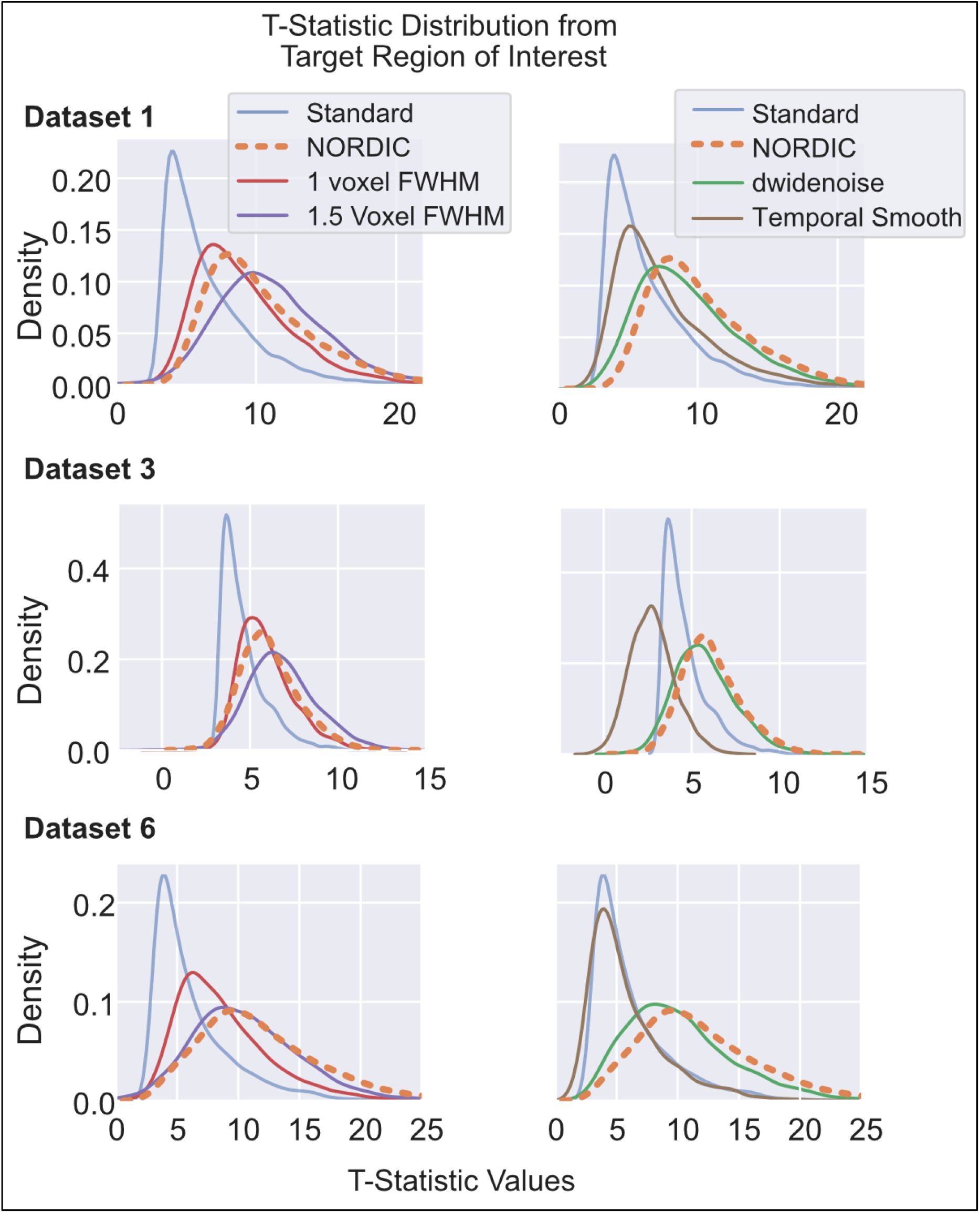
Distributions of t-statistics from alternative noise reduction methods within the Target ROI. Left column shows data from Standard, NORDIC and spatial smoothing with 1 and 1.5 voxel FWHM spatial smoothing. Right column compares the same Standard and NORDIC data against temporal smoothing and dwidenoise denoising. T-values were extracted from the Target ROI defined using the Standard data. The t-values obtained with NORDIC (Orange, dashed) processed data is comparable to the effects of an additional 1 or 1.5 voxels FWHM gaussian smoothing. While temporal smoothing (brown) did increase t-statistics for Dataset 1, note that for the fast event-related design (Dataset 3) this led to a temporal blending of neighboring events, leading to positive and negative t-values, an effect not found in NORDIC data.

Distributions for NORDIC and Standard are identical to that given in Figure 1, with the mean across all datasets at 9.9±4.5 and 6.16±3.6, respectively. The mean for t-statistics for these other methods are as follows: 9.2±4.5 for dwidenoise, 8.5±4.2 for 1 voxel of additional smoothing, 10.1±4.9 for 1.5 voxels of additional spatial smoothing, and 6.6±4.4 for temporal smoothing (Supplemental Fig. S2).

Temporal smoothing appears to confer minimal benefits with respect to (autocorrelation corrected) t-statistic distributions for the block designs used in DS1 and DS6 (Fig. 1); as can be expected, it begins to fail as a processing method when used on the fast event related design in DS3, yielding t-statistics that decrease and approach zero due to blending the events that are closely spaced in time. No such effect is seen in the NORDIC reconstruction, which is in fact right-shifted with no negative values. The performance of dwidenoise approaches NORDIC, in terms of t-statistics but, as discussed later, has a complex spatial smoothing effect on the data.

### Effects of Denoising on Global Image Smoothness

Image smoothness was estimated from the data both prior to and subsequent to processing steps that corrected for slice timing and motion (labeled as Pre- and Post-Processing in Figure 3) and on the residuals from the conventional GLM (Figure 3) (see Methods). Any spatial smoothing introduced by post-acquisition data manipulations shows up as an increase in FWHM relative to the Standard data.

**Figure 3.**
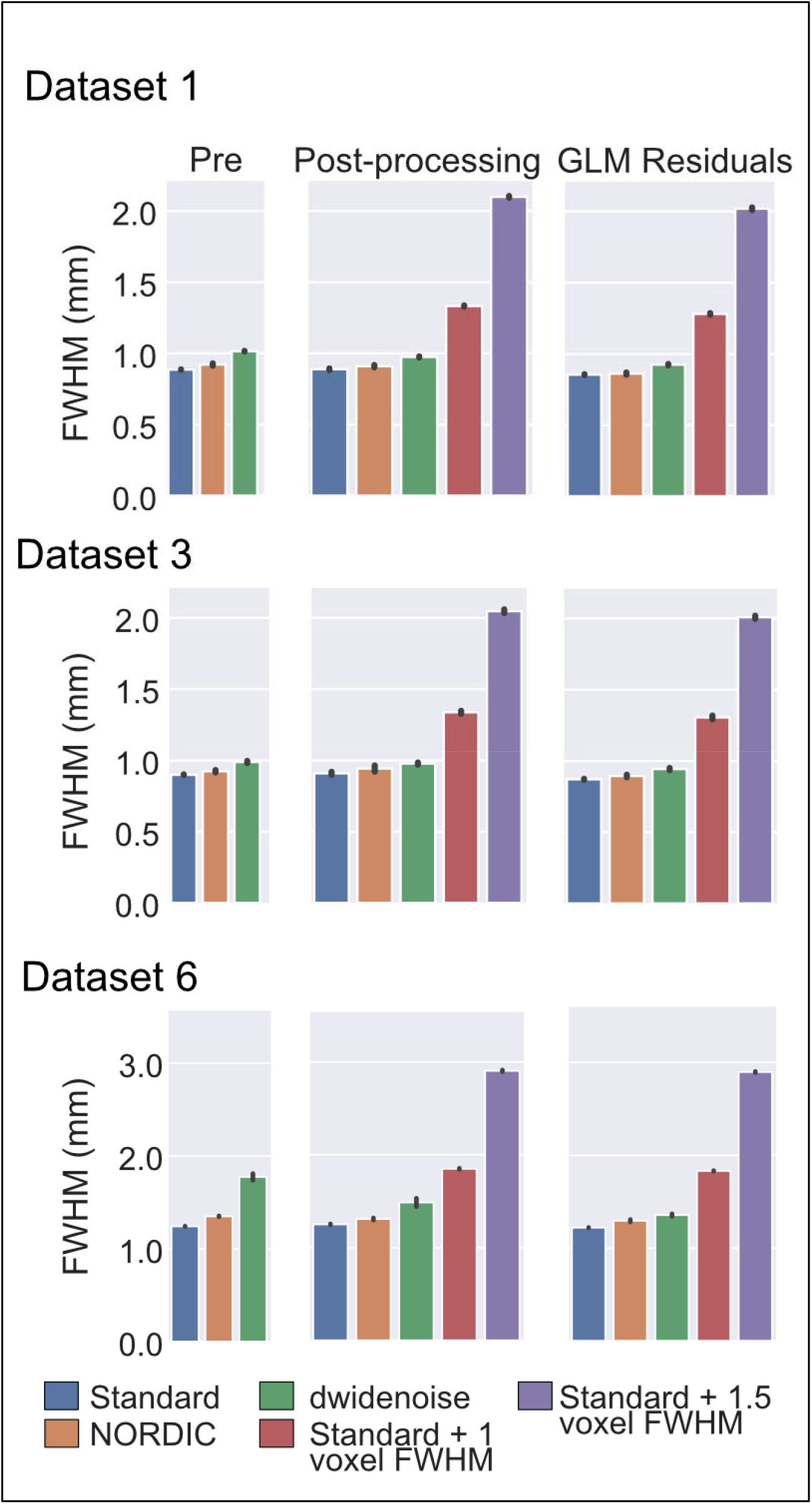
Estimated spatial smoothness in mm (FWHM) at various processing stages for each method. Prior to processing (left), NORDIC results in an average increase in 5.1% in smoothness and dwidenoise results in an increase of 22.4% on average. After processing, but prior to the GLM (middle) this trend remains. Note that the image smoothness of the Standard, NORDIC, and dwidenoise data are substantially below the level of the additional 1 or 1.5 voxels of additional smoothing. These trends remain the same for the residuals (right) after the conventional GLM. Error bars indicate standard deviation over runs.

The nominal resolution specified for image acquisition for these datasets was 0.8, 0.8, and 1.2mm isotropic, respectively. The FWHM measured in the Standard data before any processing were 0.887±0.002mm, 0.900±0.003mm, and 1.244±0.002mm, respectively, which is marginally higher than the nominal resolution specified for image acquisition. In examining the NORDIC datasets for the smoothness prior to any processing, we found a small increase in estimated smoothness associated with the NORDIC reconstruction (Table 2, Figure 3). The NORDIC data smoothness, relative to the Standard reconstruction values, corresponds to an average increase in estimated image smoothness of 5.13% across the 3 datasets shown in Figure 3. For dwidenoise, a much larger increase in smoothness was evident, with an estimated FWHM before processing corresponding to a 22% increase on average. Across all 7 Datasets (Supplemental Figure S4), NORDIC led to a 5.6% average increase in estimated image smoothness, whereas dwidenoise led to a 16% increase in image smoothness, prior to motion correction and slice timing.

**Table 2.** Estimated Smoothness in millimeters (FWHM). The first set of columns shows the estimated smoothness before any additional processing, within a brain mask made from the first run. The next set shows the estimated smoothness after motion correction and slice timing, with the bottom two rows reporting the effects of explicit, intentional smoothing. The final set of columns shows the estimated smoothness of the residuals from the conventional GLM. Across all processing timepoints the NORDIC data is minimally smoother than the Standard data. Values are mean across runs, plus/minus standard deviation.

|  | Before Processing ('Pre') |  |  | Post Processing |  |  | Residuals |  |  |
| --- | --- | --- | --- | --- | --- | --- | --- | --- | --- |
|  | DS1 | DS3 | DS6 | DS1 | DS3 | DS6 | DS1 | DS3 | DS6 |
| Standard | $0.89 \pm 0.002$ | $0.9 \pm 0.003$ | $1.24 \pm 0.002$ | $0.89 \pm 0.006$ | $0.91 \pm 0.012$ | $1.25 \pm 0.006$ | $0.85 \pm 0.004$ | $0.87 \pm 0.004$ | $1.22 \pm 0.002$ |
| NORDIC | $0.92 \pm 0.009$ | $0.92 \pm 0.01$ | $1.35 \pm 0.006$ | $0.91 \pm 0.009$ | $0.94 \pm 0.024$ | $1.31 \pm 0.012$ | $0.86 \pm 0.009$ | $0.89 \pm 0.01$ | $1.3 \pm 0.014$ |
| dwidenoise | $1.02 \pm 0.003$ | $0.99 \pm 0.009$ | $1.78 \pm 0.039$ | $0.98 \pm 0.004$ | $0.98 \pm 0.008$ | $1.49 \pm 0.048$ | $0.92 \pm 0.005$ | $0.94 \pm 0.009$ | $1.36 \pm 0.015$ |
| +1 Voxel FWHM | - | - | - | $1.33 \pm 0.005$ | $1.34 \pm 0.012$ | $1.85 \pm 0.006$ | $1.33 \pm 0.005$ | $1.3 \pm 0.015$ | $1.83 \pm 0.004$ |
| +1.5 Voxel FWHM | - | - | - | $2.1 \pm 0.007$ | $2.05 \pm 0.014$ | $2.9 \pm 0.01$ | $2.1 \pm 0.007$ | $2.01 \pm 0.012$ | $2.9 \pm 0.006$ |

Following the pre-processing steps, which were identical for all subsequent applications of “denoising”, smoothness estimates remained similar for the 3 datasets reported here (Column 2 Table 2, Figure 3). The mean increase in estimated smoothness for all 7 datasets, relative to the Standard post-processed data, was estimated to be larger by 3.3% for NORDIC, 9.3% for dwidenoise, 51% for 1 additional voxel of smoothing, and 140% for 1.5 voxels of smoothing.

Following a conventional GLM, the mean increase in estimated smoothness of the residuals for all 7 datasets, relative to the Standard post-processed data, was 3.7% for NORDIC, 8.0% for dwidenoise, 52.7% for 1 additional voxel of smoothing, and 142.8% for 1.5 additional voxels of smoothing.

For all processing stages, the increase in estimated smoothness of NORDIC was significant (all p<<0.001), as was the increase in estimated smoothness due to dwidenoise (all p<<0.001). In addition, NORDIC was significantly less smooth at all stages compared to dwidenoise processed data (all p<0.001).

### Effects of Denoising on Local Image Smoothness

Local smoothness estimates, in mm FWHM, are presented in Figure 4. We focus on the 0.8mm high resolution Datasets (DS1 – DS3) for which spatial precision is most important. The first column in Figure 4 shows the slice presented in the subsequent columns. Visual inspection of the next 3 panels shows that the local smoothness varies across the brain and tissue classes in the residuals of the task GLM; such a variation can be expected due to processes such as spatially correlated spontaneous neuronal activity (e.g. (Smith et al., 2009)) or the propagation of the high temporal fluctuations associated by veins (Chen et al., 1999; Kim et al., 1994; Zhao et al., 2006) to neighboring voxels due to the BOLD effect. These effects were minimal in the Standard data (Fig. 4A, 2^nd^ row from left), though punctate regions of high local smoothness, reminiscent of blood vessel cross sections, were visible likely as a result of the aforementioned, temporally correlated fluctuations associated with veins.

**Figure 4.**
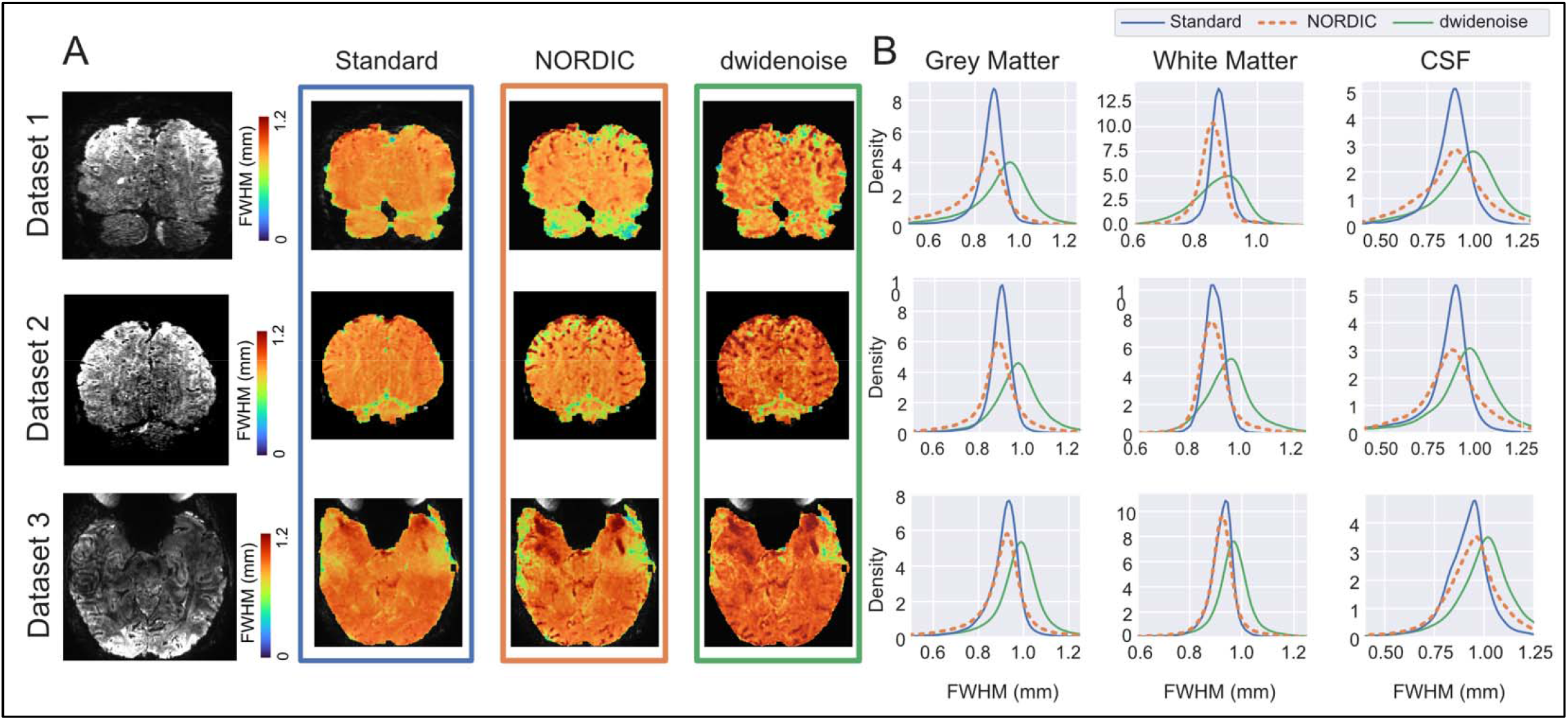
Local smoothness estimated from GLM residuals, across all runs. **A) Selected slices and local smoothness estimates, in FWHM mm.** The leftmost panel shows the selected EPI slice. The next three panels show the estimated, voxel-wise (local) spatial smoothness for the three different processing methods, Standard (blue border), NORDIC (orange) and dwidenoise (green), with the scales identical between the different processing types. Note that the local spatial smoothness is often highest in dark areas of the EPI image, likely associated with veins. **B) Full distributions of voxel-wise local smoothness estimates within different tissue classes.** These kernel density estimates show the distributions of the local spatial smoothness estimates in tissue classes derived from a T1-weighted anatomical image for Standard (blue), NORDIC (orange) and dwidenoise (green). Local smoothness is somewhat decreased following NORDIC, except within the CSF mask. All datasets had a prescribed resolution of 0.8mm isotropic.

Following NORDIC processing, voxels within regions corresponding to white matter have similar or reduced spatial correlation of temporal signatures (Fig. 4A and 4B), whereas gray matter is more variable across these presented datasets; the punctate regions present in the Standard are now more clearly visible (Fig.5A, 3^rd^ row from left). Following dwidenoise, there is a general increase in the local FWHM estimates across the entire brain (Fig.5A, 4^th^ row from left, and Fig. 4B). The distributions of the local smoothness estimates for all voxels within each tissue class from segmentation are shown in Figure 4B. Across the three high resolution (0.8mm isotropic) datasets shown, the mean FWHM with the gray matter mask was 0.87±0.02mm for Standard, 0.87±0.05mm for NORDIC, and 0.96±0.0mm for dwidenoise. The mean FWHM in white matter was 0.90±0.02mm for Standard, 0.89±0.04mm for NORDIC and 0.93±0.06mm for dwidenoise. The mean FWHM in CSF was 0.88±0.04mm for Standard, 0.91±0.04mm for NORDIC and 0.98±0.03mm for dwidenoise (see Supplemental Table S1 for individual dataset values).

**Figure 5.**
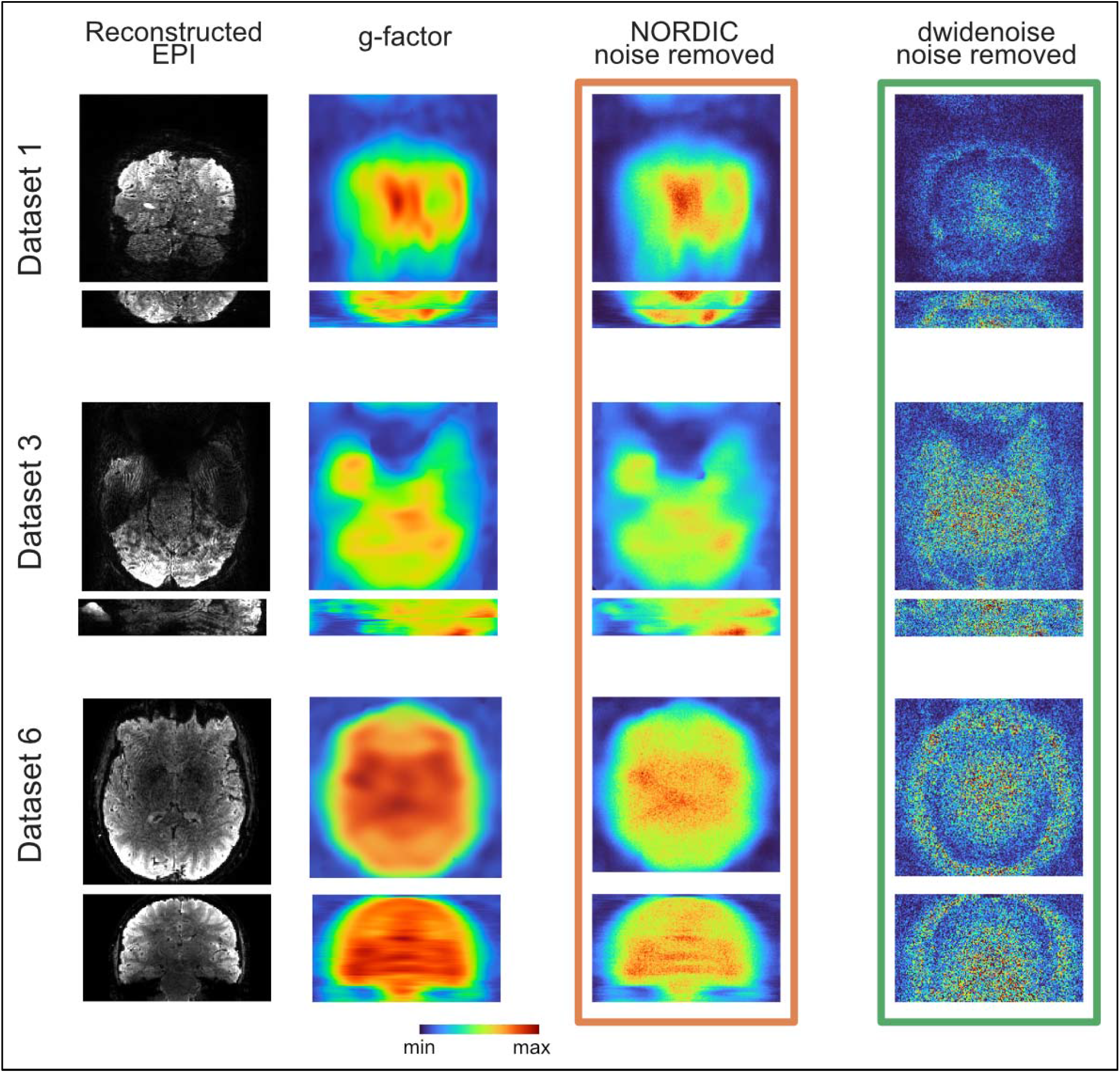
Comparing Residuals from Denoising Methods. The first column shows the selected views using the reconstructed EPI images. The next column shows the g-factor maps calculated from the raw k-space data. The last two columns show the temporal mean (absolute values) of the extracted noise from the first run for NORDIC (column 3) and dwidenoise (column 4).

Across all 7 datasets, relative to the Standard data, the mean local smoothness estimates for NORDIC were 2.9% smaller in gray matter, 9.5% smaller in white matter and 11.7% larger in CSF (all p<0.001). For dwidenoise, smoothness estimates were 2.8% greater in gray matter, 7.9% smaller in white matter and 14.3% larger in CSF (all p<0.001). For all tissue classes, NORDIC had significantly lower estimates of local smoothness relative to dwidenoise (all p<0.001).

### Spatial Characteristics of Extracted Noise

Following image reconstruction and denoising, we first examined the spatial characteristics of the noise amplification due to accelerated image acquisition (’g-factor’) as well as the noise removed from the data timeseries by NORDIC and dwidenoise. Figure 5 shows the images for 3 data sets comparing the temporal mean of the extracted noise from the first run with the maps of the g-factor produced from the raw k-space data. The g-factor maps essentially reflect the spatial distribution of the thermal noise component in the data with the spatially non-uniform amplification that comes from the use of parallel imaging. NORDIC processed data demonstrate that the image of what is removed looks similar to the g-factor map, without any hint of brain related structures or edges. In contrast, anatomical boundaries are visible in the dwidenoise data.

### Evaluating temporal precision

#### Fourier Analysis

The normalized power spectra for DS7 and DS8 show that denoising by NORDIC results in a wide reduction in power across frequency bands, with the effects most apparent at higher frequencies where thermal noise is the dominant noise source (Fig. 6). This effect is most visible in DS8, which was collected at a TR of 350ms. At this sampling rate, the frequency associated with cardiac noise (∼1Hz) is clearly visible in the data and clearer after NORDIC processing. The task related frequency (0.05Hz) is also more pronounced after NORDIC for DS8. Though this effect is most visible in the gray matter partition, it is also visible in the white matter. This is likely a consequence of both partial volume effects and imperfect ROI overlap between the anatomically derived tissue segmentation and the distorted EPI images.

**Figure 6.**
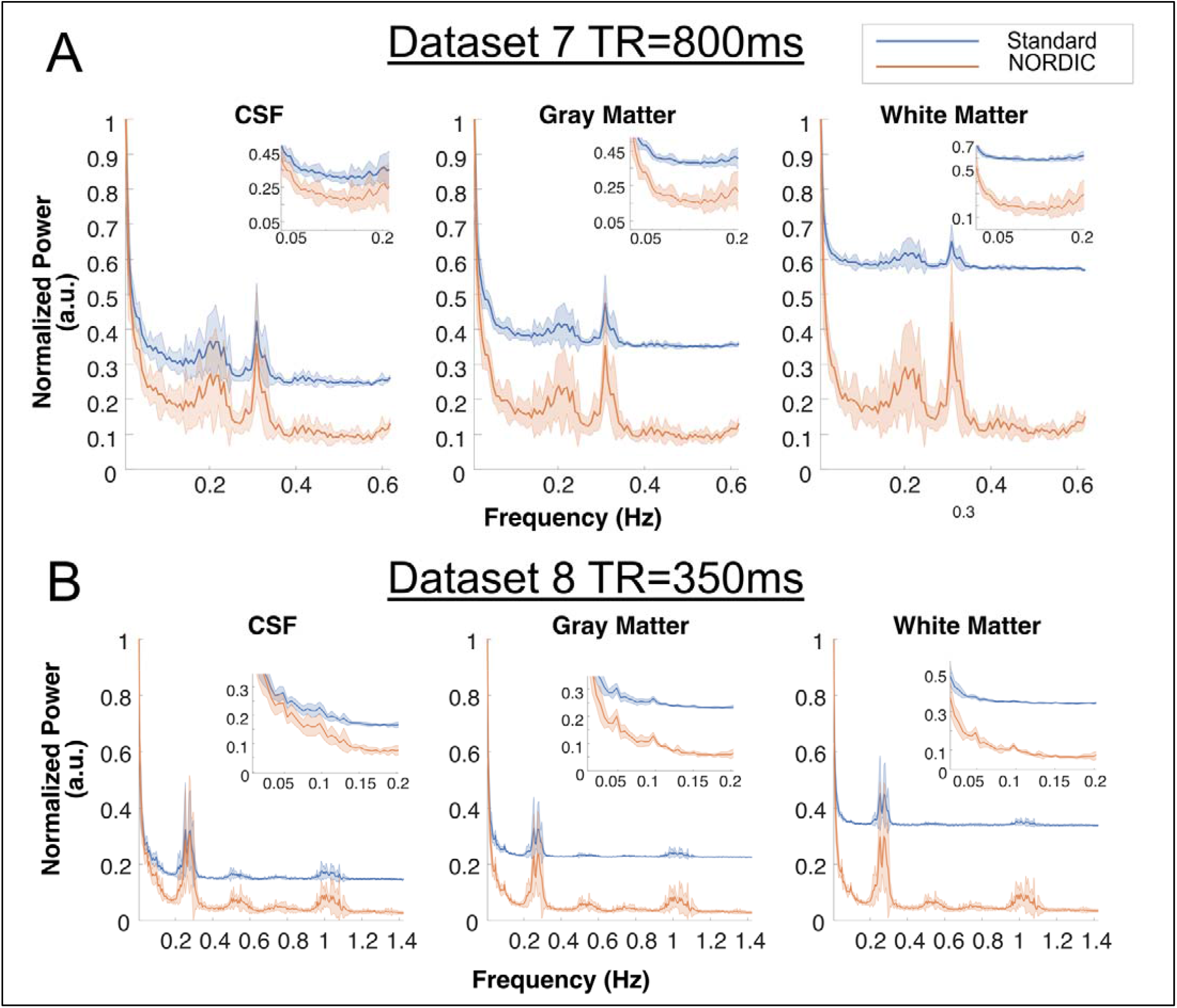
**Frequency Plots from FFT analysis**. **Panel A)** Normalized power spectra from DS7, a 3T, HCP-like acquisition with 800ms TR. NORDIC processing reduces power throughout most frequency bands, with effects such as a task harmonic (∼0.2 Hz) and the respiratory band (∼0.3 Hz) becoming more visible. Insets show power from 0.01 through 0.2. **Panel B)** Normalized power spectra from DS8, a rapidly sampled acquisition with 350ms TR. The effect of NORDIC in broadly reducing power remains pronounced throughout higher frequencies. Here the respiratory and cardiac signals are clear at ∼0.3 and ∼1 Hz respectively. In the gray matter, a clear peak at 0.05Hz, corresponding the task frequency is also clearer after NORDIC processing (see inset). Shading shows standard deviation across independent runs for both A and B.

#### Cross Validation

Following NORDIC processing, estimates of single-run and single voxel HRFs from the FIR model were markedly improved. We find that following processing with NORDIC, the FIR time courses are more consistent with typical hemodynamic-like responses (i.e., approximate a double gamma) and display, on average, 33% less variability across runs (Supplemental Table 3 shows average variability for each dataset within the Target ROI). To see the origin of these improvements, we consider 81 voxels selected from DS1 focusing on an area that features both target and non-target sensitive voxels. Figure 7A (top row) shows the underlying data and activation maps from the conventional GLM across all runs of the Standard processed data, highlighting in the inset containing the 81 voxels considered for Figures 7B and 7C. Visual inspection of individual voxel time courses (with activation map overlaid) from a single run (the first run) of DS1 in Figure 7B shows that the reduction in noise from the NORDIC method (right panel) does not lead to a spread of the activation (consistent with the negligible change in spatial smoothness) but instead reduces the noise level such that stimulus-coupled signal changes becomes more visible. The selected 81 voxels contain responses to the target (indicated by #1, #2), responses to target and surround (within grey boundary), and responses only to the surround (indicated by #3). Despite identical voxels being selected for the Standard data (left panel), stimulus-evoked responses are difficult to see. Though the spatial maps presented in Figure 7A (upper row) was derived from the full 8-runs, the task events are visible in the individual voxels of the single run after NORDIC processing (Fig. 7B, right panel) but generally not in the Standard. Section C shows response estimates from an finite impulse response (FIR) model for the selected voxels, showcasing improvements in single-run, single voxel FIR estimates.

**Figure 7.**
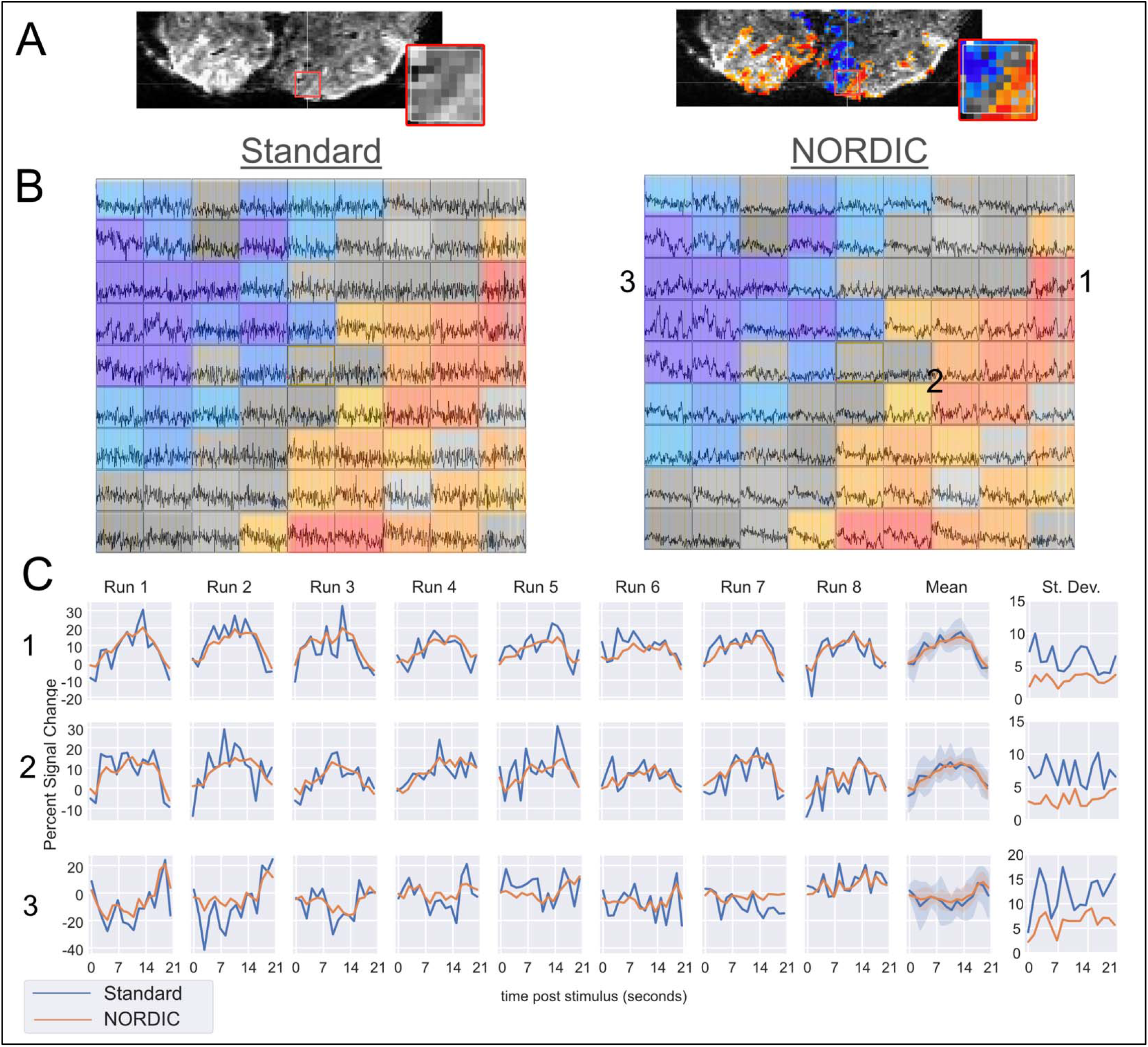
Example of activation, improved time courses and FIR estimates in run 1 of DS1, Panel A, upper row Different views of the area under consideration from Standard reconstruction: Left column shows the mean GE-EPI values for the area under consideration. Right shows the activation amplitude (−12 to 12 % signal change) in the selected slice for all runs, Center > Surround from the Standard data (map threshold from t-stat ≥ 3.3). Inset boxes in panel A show the 81 voxels considered in Panel B. **B) Time courses from first run for 81 voxels.** To visualize task responsive voxels, we shade them based on the contrast from all runs of Standard data, as seen in Panel A, right. The stimulus-evoked signal amplitude changes associated with the three surround and the three target stimulus epochs are clearly visible in the NORDIC processed (Right) timeseries of the corresponding voxels but are largely invisible in the Standard (Left) data due to high noise levels. **C) Single Run FIR Estimates for the Target Condition.** Responses to the target (center) are illustrated in selected voxels 1 and 2 for individual runs are shown. The final columns show the across-run average and standard deviation respectively. Shading in the across-run average plot shows standard deviation from the mean, which is also plotted separately for clarity (note that the deviation associated with the Standard data far exceeds that of NORDIC). Voxel 3, which is sensitive to the surround condition, remains closer to the expected zero amplitude (i.e., non-responsive). This is particularly true for NORDIC processed data, which is associated with lower standard deviation.

To further investigate whether NORDIC processing does indeed preserve the inherent signal present in the fMRI data (i.e., only suppressing thermal noise), we considered an exhaustive Leave-p- Out cross validation scheme to determine how effectively FIR model results could predict held-out timeseries quantified with the coefficient of determination, R^2^ (See Methods).

Panel A in Figure 8 shows the performance of a *single* run of Standard and NORDIC data in predicting the full timeseries of held out data. In general, the R^2^ metric increases as voxels with better predicative accuracy in the full model are included, with all lines tending to increase from left to right. Notably, however, a single run of NORDIC is better able to predict the held out runs of Standard data compared to the Standard data itself. Further, this benefit is maintained even for voxels that had a high signal to noise ratio (i.e., far right of the graph). Exemplar estimates from the finite impulse response (FIR) model which produces an estimate of the HRF are shown for a single voxel from a single run in the lower right of each graph in Figure 8A. As expected for such high-resolution data, the estimates from the Standard data (blue) are noisy. However, following NORDIC processing, these single run estimates show clear HRF-like properties.

**Figure 8.**
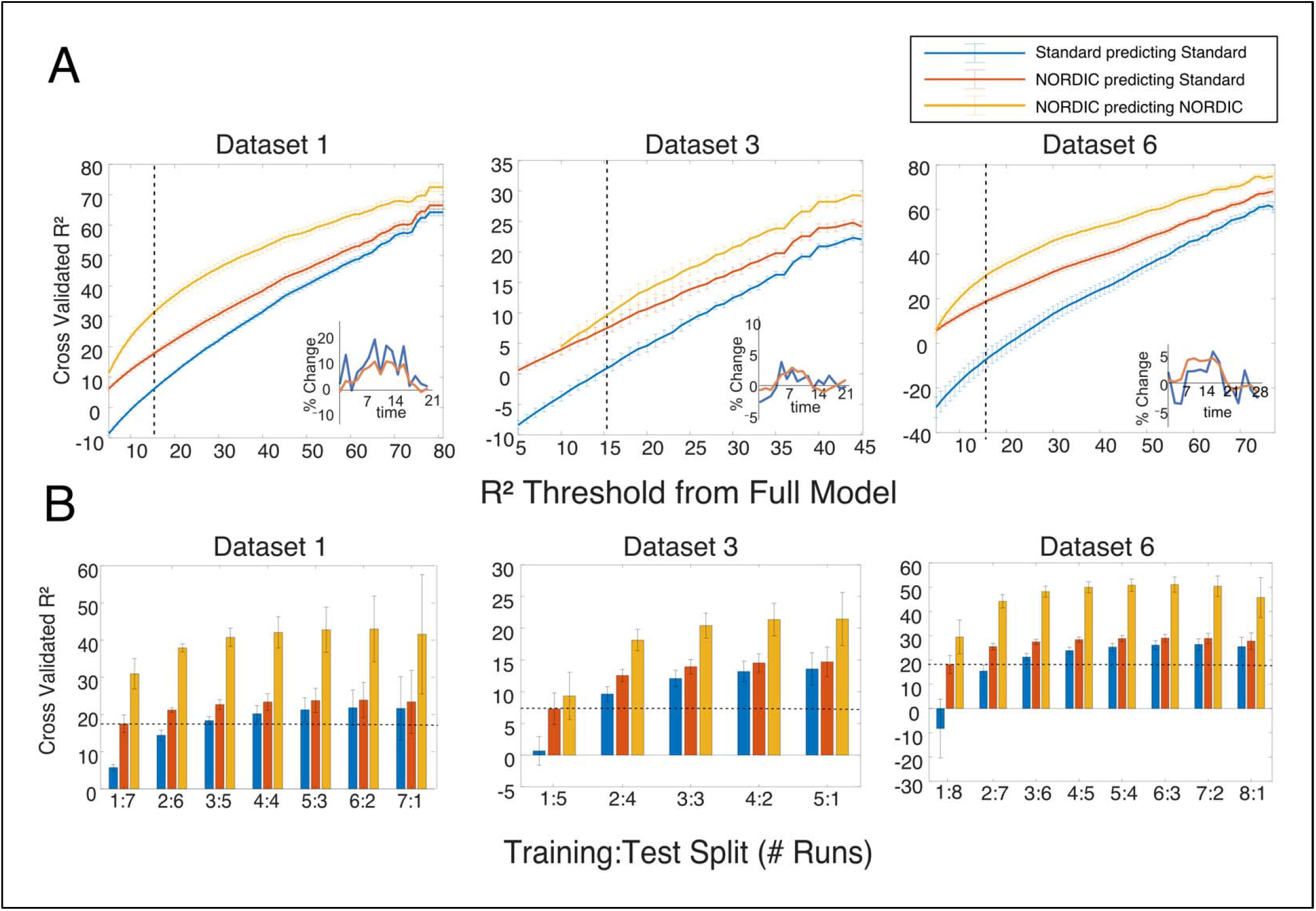
NORDIC processing leads to higher cross-validation performance, predicting from a finite impulse response model. **Panel A)** Exhaustive cross validation performance when training on using only one run. Cross validated R^2^ is shown for training on Standard data and predicting held-out Standard data (blue), training on NORDIC data and predicting held-out NORDIC data (yellow), and training on NORDIC data and predicting held-out Standard data (Orange). X-Axis indicates voxel inclusion threshold derived from leave-one-out cross-validated R^2^ using a canonical HRF on Standard data. Insets show example single voxel single run FIR model estimates for Standard (blue) and NORDIC (Orange). NORDIC processing can produce estimates that better predict Standard data compared to Standard data itself. Error bars are standard error over permutations. Dashed lines show an R^2^ threshold of 15% used in panel B. **Panel B)** Leave-p-Out training was repeated for all Ps less than the number of runs, N-1. Colors are as above; the number of runs included in the training vary across the X-axis, with bar height reflecting the R^2^ obtained. Dashed line indicates the performance of training on one run of NORDIC data, which is equivalent in cross validation performance to using 2 or 3 runs of Standard data. Including more data allows Standard models to approach, but not reach 2 to 3 runs of NORDIC data. Error bars again show standard error across permutations. Error bars in A indicate standard error and those in B indicate standard deviation.

To quantify this improvement across all voxels and runs we varied the number of testing runs, P, from 2 to the number of runs-1 (Figure 8B). Using a threshold of voxels that were able to explain 15% of the variance in the full model (vertical lines in panel A), we can see that training with one run of NORDIC is able to predict a held out timeseries as well as 2 to 3 runs of Standard data (horizontal dashed line, Panel B), and two runs of NORDIC are nearly able to predict as well as any number of Standard runs combined. Across all thresholds and folds, NORDIC processed data is also always able to better predict the timeseries of data that has undergone NORDIC processing - which would be the typical use case. These findings apply to all datasets considered (Supplemental Figure S5).

## Discussion

Most applications of the MRI method are highly SNR limited. As such, it is common to perform some sort of noise mitigation procedure in post-acquisition data processing in order to increase the SNR of the data. Although the ultimate goal is to do so without compromising any information, conventional methods produce a trade-off, such as reducing effective spatial and/or temporal precision. For example, spatial smoothing, an extremely common strategy, achieves SNR gains by averaging the data over spatial coordinates of the image, thus inducing blurring. More contemporary denoising methods (e.g. (Pruim et al., 2015; Thomas et al., 2002; Veraart et al., 2016), and references therein) try to characterize the components of the data and selectively remove some of them. As it is always possible that the effects of denoising can be deleterious, is imperative that a careful and critical evaluation is performed when deploying such an approach, especially when substantial and potentially transformative gains are promised for the field of interest, as is the case with the application of NORDIC to fMRI (Vizioli et al., 2021).

In this paper, we extend an evaluation of the recently described NORDIC denoising method as applied to fMRI to a wider variety of field strengths, voxel sizes, TRs, and stimulus designs. We find that NORDIC-processed fMRI data removes noise that matches g-factor maps (Fig. 5), producing much higher t-values under a conventional fMRI modeling framework (Figs.1, 2), consistent with the prior report (Vizioli et al., 2021). The fMRI t-statistics achieved after NORDIC denoising are approximately equivalent to those produced by smoothing the data using a kernel of 1.5 voxels; however, analysis of the NORDIC-processed data does not show any comparable increase in smoothness (Figs. 3,4). The frequency spectrum of data following NORDIC shows a widespread reduction, consistent with white-noise suppression (Fig. 6). These SNR gains are also reflected in markedly improved single run, single voxel FIR estimates, which in turn produce better predictions of held-out, non-denoised Standard data (Fig. 7,8), compared to the Standard data itself. Collectively these findings support the use of NORDIC for a wide variety of fMRI applications, ranging from HCP-like data obtained at 3T to cutting edge high-resolution fMRI data acquired at ultrahigh magnetic fields.

### Conventional GLMs

The t-statistic is widely used in the fMRI literature to report the statistical veracity of signal increases and decreases as spatial maps. Consistent with our prior report (Vizioli et al., 2021), we find that NORDIC-processed data had substantially larger t-values compared to the Standard data alone (Figure 1). These gains are similar to or greater in magnitude than those reported by others using denoising methods to remove *structured* noise such as multi-echo denoising (Gonzalez-Castillo et al., 2016; Kundu et al., 2017), ICA-based denoising strategies such as ICA-AROMA (Pruim et al., 2015) or using SNR efficient accelerated imaging sequences like SMS/MB (Moeller et al., 2010) to collect more data in a given period of time (Smith et al., 2013). However, NORDIC is a complement rather than a replacement to these methods as it focuses on suppressing thermal noise. We observe that this gain is not due to a large shift in the estimated activation amplitude, as the betas remain highly similar following NORDIC processing (Supplemental Figure S6).

Of the other processing methods examined here, the performance of NORDIC with respect to t-statistics exceeds all but the 1.5 voxel spatial smoothing (Figure 2), even in the data with relatively large voxels (i.e., 2mm isotropic resolution 3T HCP protocol). Of course, as previously mentioned, NORDIC accomplishes this without meaningful increases in estimates of blurring. NORDIC also outperformed the benefits one would get with temporal smoothing, even in cases where long duration (i.e., 12s) events were separated by long inter-stimulus intervals. While NORDIC does produce a timeseries that is less corrupted by thermal noise, we did not detect effects that would be consistent with averaging over a temporal window. This is most clear in DS3, which used a fast event related design. Following temporal smoothing, the t-values for the face condition in this design decreased. This reflects the mixing of neighboring events due to the short ISI of 2 seconds. The opposite effect is found following NORDIC processing, in other words, t-values increased, and negative t-values were absent. Under this approach, we did not observe that temporal precision was lost after NORDIC processing. Both spatial and temporal smoothing also, as expected, additionally alter activation amplitudes an effect which is not observed on the NORDIC processed data (Supplemental Figure S6).

These findings present the possibility of new avenues of research, ultrahigh spatial and/or temporal resolution studies, the use of smaller ROIs for ROI-based analysis, or examining single-trial response estimates (Chen et al., 2021), all of which represent important but often SNR-starved analysis strategies.

### Image Smoothness

Despite the similarity of the distributions of the t-values between NORDIC and spatially smoothed data, the NORDIC data is not associated with a comparable increase in the estimated spatial smoothness (e.g., +1.5 voxel FWHM smoothness estimated to be 132% larger; Figure 3). In fact, at its maximum, NORDIC only increased the estimated smoothness by 6.1%. This is smaller than the effect often observed with conventional preprocessing methods, which are known to produce images with greater spatial smoothness characteristics due to the need to interpolate values on a new image grid (Polimeni et al., 2018). While these effects were significant, they were very small (more than 1 or 2 orders of magnitude less than 1 or 1.5 voxels of spatial smoothing respectively) and did not compromise cross-validation accuracy (Figure 8, S5), nor do they match the effects of spatial smoothing when comparing betas (Figure S6).

Here we considered estimates of global smoothness at all stages of data processing in the fMRI data analysis (Figure 3). While typical smoothing estimates use the residuals of the data as an estimate of the overall smoothness of the noise, it is possible for these estimates to be overestimated. For example, coherent areas of signal change could remain due to a mismatch between the canonical HRF and the subject’s response. The patterns of smoothness reported here are consistent among processing stages, supporting the argument for minimal image smoothing due to the NORDIC method. This global measure of image smoothness is often used in the context of cluster correction, as it is thought to reflect the underlying smoothness of the acquired image (Cox et al., 2017), but has also been used to evaluate processing and acquisition approaches (Esteban et al., 2019; Friedman et al., 2008, 2006; Marcus et al., 2013).

This finding is further corroborated when we examine the spatial autocorrelation of each voxel’s correlations with its neighbors, which we have termed ‘local smoothness’. NORDIC processing produced local smoothness estimates that were nearly equivalent to the standard data (Figure 4). We did observe a small decrease in the estimated local smoothness following NORDIC processing for gray and white matter. As this metric is computed on the residuals of the GLM, it is plausible that the model obtained a better fit for task responses or structured noise, such as motion, after NORDIC processing. As such, this is not reflecting an increase in the spatial resolution of data following NORDIC processing, but instead likely highlights that the model captured more of the structured variance in the signal.

We observed a larger positive deviation within the CSF mask, which includes features such the superior sagittal sinus as well as punctate regions likely associated with cross sections of blood vessels, which, in case of veins, appear as also dark punctate structures in the anatomical images (Figure 4a, left most column). Macroscopic blood vessels large enough to be seen in these images are expected to have relatively large signal fluctuations, as was shown for veins in previous fMRI studies (Chen et al., 1999; Kim et al., 1994; Zhao et al., 2006). These fluctuations exist independent of the stimulus or task in an fMRI experiment. Especially in case of the veins, these fluctuations will extend beyond the boundaries of the blood vessel into neighboring voxels due to the BOLD effect. Such correlations are expected to be “unmasked” and easier to detect after the suppression of thermal noise, leading to an increase in the size of the region of locally smooth, correlated voxels. Similarly, when the thermal noise is suppressed by NORDIC, it unmasks higher local correlation due to the BOLD effect associated with pial veins as well as to other physiological processes such as pulsations due to heartbeat and respiration in the CSF space.

These local smoothness findings are different from the global smoothness estimates, in that, on average, NORDIC processed data had marginally less estimated smoothness in both gray and white matter. One possible source of this difference is likely due to the difference in analytical methods. For example, global smoothness estimates consider each volume independently, in effect examining the variance across space. In contrast, local smoothness considers autocorrelation of each voxel’s timeseries correlation within a local neighborhood. It is plausible that neighboring voxels could have highly correlated timeseries, despite large differences in signal magnitude (i.e., high spatial variance), such as at gray/white matter boundaries due to partial volume effects. It is also possible that the global smoothness after NORDIC seemed slightly increased in the GM due to effects similar to those described above for the CSF mask. Nevertheless, both global and local smoothness estimates provide evidence that NORDIC processing is not leading to meaningful increases in smoothness. The spatial autocorrelation methods (both global and local) to estimate smoothness in this work are different from approaches that estimate a functional point spread function (Shmuel et al., 2007), which instead attempts to quantify the functional precision available in the maps of functional responses. For the latter, prior work found that NORDIC had no impact on the functional point spread of the BOLD signal (Vizioli et al., 2021). To further validate these results we performed an initial evaluation using the local perturbation response (LPR) method (Chan and Haldar, 2021) and were able to recover the injected synthetic sparse signal, though sufficiently low intensity perturbations (i.e. below or near thermal noise level) were not perfectly recovered (See Supplement, Figures S8, S13). Further work is required to determine interpreting these results, as with all synthetic manipulations, it is difficult to match all of the properties of the natural fMRI signal.

Together, these reports show that NORDIC is able to suppress thermal noise at a level similar to that of 1 or 1.5 voxels of smoothing but avoids the increases in spatial autocorrelation associated with such levels of blurring, and instead only marginally affects the spatial properties of the signal.

### Temporal Precision

The use of NORDIC produced fMRI voxel time courses in which responses to task events were more visible and not subject to any apparent smoothing (Figure 7). We first examined the normalized power spectrum of DS7 (800ms TR) and DS8 (350ms TR). For both datasets, task and physiologically related frequencies are clearer after NORDIC processing, relative to the Standard data (Fig. 6). In general, there is a large reduction of power at nearly all frequencies, consistent with the reduction of normally distributed noise. This corresponds to an increased ability to identify and resolve frequencies associated with physiological noise in each individual voxel timeseries, however the potential utility of this for physiological denoising was not explored.

We then used the estimates of the hemodynamic response function (HRF), produced by finite impulse response (FIR) models, to simultaneously examine the denoising performance of NORDIC and whether this resulted in a substantial (i.e. affecting cross-run accuracy) loss of temporal information. NORDIC produced FIR estimates that were associated with less cross-run variability (Figure 7, Supplemental Table 3). In many cases, particularly in high-resolution studies, these types of response estimates are produced by simultaneously modeling multiple runs or averaging the signal within an ROI. Here, however, we show that single-run estimates are reliable, even at the single-voxel level.

The primary concern is that these estimates are the result of suppressing both signal and noise. That is, the process of removing thermal noise has also removed signal sufficient to alter the measurable temporal information in the fMRI time course. We do not observe this effect in the NORDIC data as shown by the fact that these FIR estimates accurately reconstruct data that were held-out from the model. This was performed in an exhaustive Leave-p-Out fashion, considering all combinations of 1≤P<Number of runs. Based on the coefficient of determination (R^2^), not only was NORDIC data better able to predict held out NORDIC data, but that it was also better able to predict held-out Standard data (Figure 8). This is a critical feature in considering the performance of a denoising method and, of course, is not always achieved. For example, large amounts of spatial smoothing will lead to increased t-values, but the voxel-wise HRFs derived from smoothed data will no longer correspond to the precise spatial location of voxels in unsmoothed data. Additionally, these activation amplitudes will have altered magnitudes (Supplemental Figure S6).

Across all datasets, the NORDIC data were better able to predict the Standard data as measured by the coefficient of determination, including datasets using acquisition methods for which the thermal noise contribution is lower (i.e., datasets with larger voxels) relative to physiological fluctuations. In order to further examine the possibility of a loss of temporal precision we also probed the neighboring timepoints in the previously mentioned LPR analysis (Chan and Haldar, 2021) and while this sparse (i.e. does not repeat over the timeseries) and synthetic signal is measurable at subsequent timepoints following denoising with NORDIC (Figure S8), the artifact was nearly 2 orders of magnitude smaller than intrinsic timeseries fluctuations (Figure S11) and as such, is effectively invisible in voxel time courses (S10).

The NORDIC processed data were also able to better predict NORDIC timeseries (Figure 8). While less critical than the above demonstration of signal preservation, this indicates that the effects of NORDIC are consistent from run to run. In this context, one (DS1, DS6) or two (DS3) runs of NORDIC have better cross validated performance using voxels that survive the 15% R^2^ threshold (Figure 8) than any number runs of Standard data. As the typical fMRI experiment would employ similarly denoised data throughout all analyses, rather than testing against the standard data (as was done here for validation), these large SNR gains represent the expected benefit of using NORDIC. Since NORDIC denoising is done for each run separately, the data from separate runs remains statistically independent. This, in conjunction with the large gains in cross validated performance may allow analyses approaches which previously required large regions of interest to be performed on the level of individual voxels. Furthermore, these improvements could translate to shorter scanning times, with many added advantages, for example, decreasing the possibility of motion and time burden for participants or patients.

NORDIC is expected to complement denoising strategies that remove structure noise, such as ICA-AROMA (Pruim et al., 2015), multi-echo ICA denoising with tedana (DuPre et al., 2021; Kundu et al., 2017) or those that leverage multiple runs, such as GLMDenoise (Kay et al., 2013); however experimental demonstration of this remains to be performed.

### Comparison with Alternative Methods

Widely used methods to remove thermal noise have only recently developed and evaluated for diffusion MRI. In functional imaging, the growing interest in higher and higher resolutions and the capability of collecting such data with reasonable sampling rates has led to increased attention to the thermal noise contribution. While thermal noise is not typically the dominant noise source in most fMRI studies (Triantafyllou et al., 2011), thermal noise begins to dominate with voxel volumes below approximately 3mm isotropic at 7T and therefore substantially impede accurate detection of signals of interest.

Choices for the reduction of thermal noise are limited, and functional neuroimaging has primarily depended on temporal averaging or spatial smoothing. Averaging requires large time commitments and can be complicated by difficulty in aligning across multiple runs, sessions or participants, while spatial smoothing with gaussian kernels unavoidably leads to a loss in spatial precision which is often the expressed purpose of high-resolution fMRI. While more advanced smoothing methods have been developed, which constrain smoothing on the basis of anatomy (Blazejewska et al., 2019; Huber et al., 2021), these methods are associated with a tradeoff – for example averaging across cortical depth may allow for high resolution analyses across the cortical surface, but necessitates the loss of depth dependent activity profiles, which are not uniform.

An alternate PCA based denoising method considered in this manuscript, dwidenoise, was developed primarily to suppress thermal noise in diffusion imaging (Cordero-Grande et al., 2019; Veraart et al., 2016); it has recently been used for resting state fMRI (Adhikari et al., 2019) and fMRI for evaluate for presurgical mapping (Ades-Aron et al., 2021, p.). However, these studies lacked a detailed analysis of the impact and the generalizability of denoising on the fMRI data, and critically, did not examine higher (sub-millimeter) resolution fMRI where thermal noise dominates. Here we find that dwidenoise does offer large improvements in typical task-based activation measures such as t-statistics; however, this appears to be at the cost of increases in estimated image smoothness. This is most apparent for the high-resolution 7T (0.8mm; DS 1, 2, 3) and 3T (1.2mm; DS6) datasets, for which precision is most desired (Figure 3). Most importantly, an image of the components removed by dwidenoise demonstrate the presence of structures that correspond to the anatomy of the imaged object, indicating that components removed are not just thermal noise. This is consistent with the suggestion that it is difficult to precisely identify the components that are removed in the MPPCA/dwidenoise approach, although its application leads to apparently better results (Moeller et al., 2021).

While these conclusions hold for our usage of dwidenoise in the present work, it is entirely plausible that further improvements could be achieved by manipulating various elements of the dwidenoise implementation. For example, it is plausible that the default settings of dwidenoise which were validated on diffusion imaging data should be altered when applying to fMRI images. In addition, it is possible to apply dwidenoise (at least for diffusion data) in complex space (Cordero-Grande et al 2019). While this or other manipulations of dwidenoise for fMRI were not tested in the current work, it is possible that this would lead to improvements in the performance of dwidenoise.

### Limitations

Although a large variety of datasets were considered in this work, including different TRs, voxel sizes, event designs, stimulus categories, and field strengths, the present work only evaluated gradient echo BOLD functional imaging, by far the most commonly employed strategy for functional imaging. The principles of NORDIC are expected to work equally well with other approaches of functional mapping, such as spin echo (SE) based BOLD fMRI (e.g. (Yacoub et al., 2003)), or functional mapping based on non-BOLD contrast mechanisms such as blood flow changes (e.g. ASL (Roberts et al., 1994) and VASO (Huber et al., 2018)). NORDIC will likely be more useful for these other functional imaging approaches since they inherently have poor sensitivity and, at any given spatial resolution, will be more limited by thermal noise associated with the MR measurement compared to GE BOLD fMRI.

Our current work suggests that NORDIC can be viewed as another processing step in fMRI which is associated with measurable, but small changes in data parameters. As such, researchers should inspect their data following NORDIC to ensure that the data is not adversely affected, particular in low SNR areas or task designs. In the datasets considered for this manuscript, the gains of NORDIC were achieved with minimal impacts on estimated image smoothness. Here we used estimates of image smoothness over global and local scales. While such FWHM measures are widely used, are sensitive to the application of image smoothing and agree with our findings in prior work which examined the functional point spread (Vizioli et al 2021), it is possible that they do not capture all of the effects of NORDIC processing. Likewise, it is possible that some temporal information is lost, however, we did not detect any negative effects in the cross-validation approach used here, and additionally observed that the fMRI signals of interest following NORDIC processing were more similar from run to run.

While NORDIC was highly effective in the data shown here, further work evaluating the effects of NORDIC, particularly for other fMRI sequences and a larger array of brain areas, is needed.

## Conclusion

The NORDIC method is suitable for use across a diverse array of functional imaging acquisition strategies in order to decrease the contribution of thermal noise. Processing data with NORDIC consistently results in substantial gains in t-values, such as those seen following smoothing, without a comparable or even moderate increase in estimates of image smoothness. In addition, NORDIC preserves the voxel-wise temporal information and is better able to predict held out data. These findings support the use of NORDIC to increase the functional contrast-to-noise ratio of fMRI, thereby improving HRF estimates and/or permitting reduced fMRI acquisition times – potentially enabling entirely new study designs and statistical approaches to data analysis. These attributes are of particular importance for ultra-high spatial resolution functional neuroimaging data targeting mesoscopic scale organizations, which are SNR-starved even at ultrahigh magnetic fields and even after extremely long data acquisitions. Similarly, acquisitions that use high temporal sampling rate of the fMRI time course, as desired for example in resting state fMRI, are also SNR starved in the individual images acquired should benefit from NORDIC substantially.

## Acknowledgements and Funding Sources

The authors would like to thank Dr. Kendrick Kay for his input. This work was supported by NIH grants: U01 EB025144 (K.U.), P41 EB027061 (K.U.), P30 NS076408 (K.U.) and RF1 MH116978 (E.Y.), and RF1 MH117015 (G.G.)

## Supplemental Material

**Figure S1.**
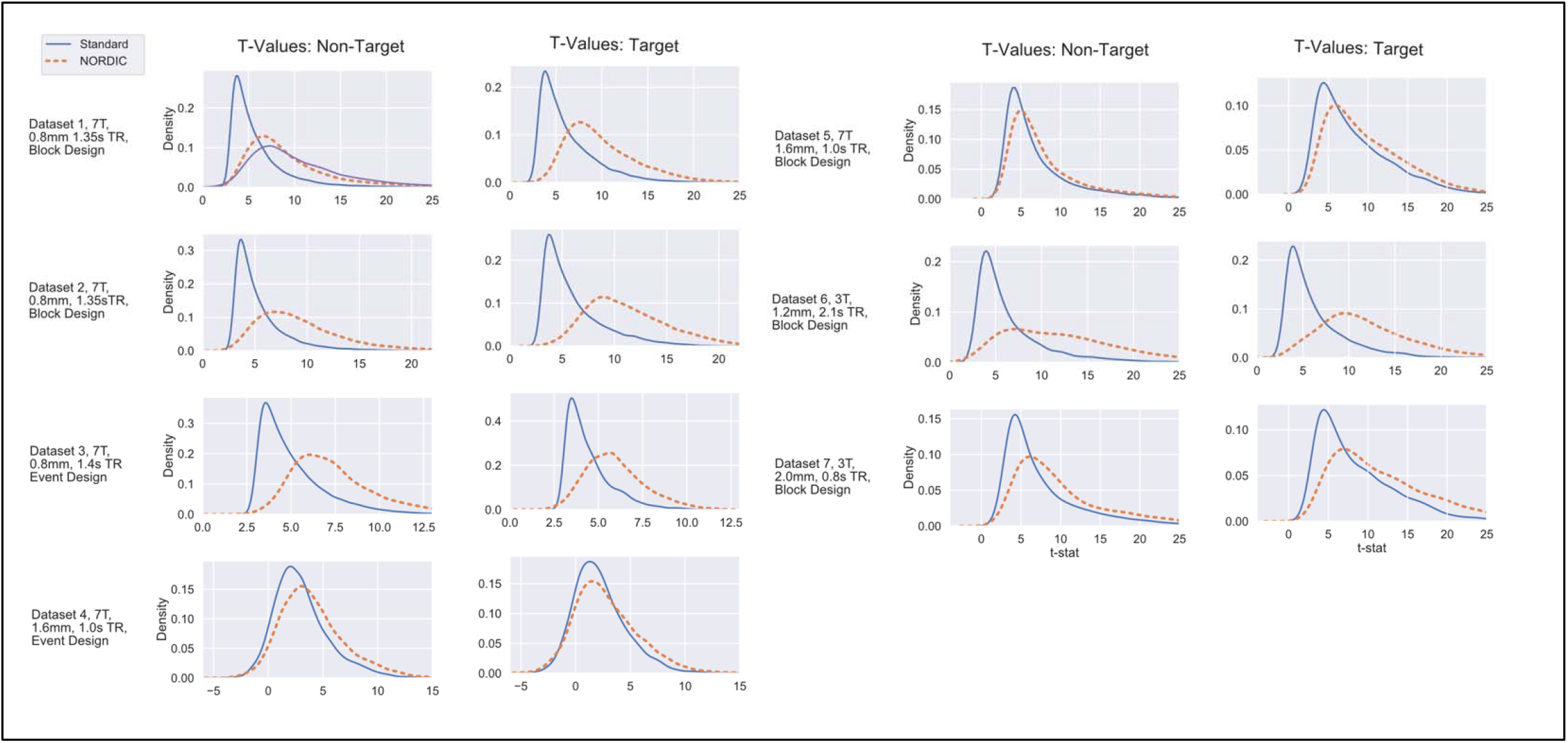
T-statistic histograms comparing Standard and NORDIC data within the Non-Target and Target ROIs. The distributions of t-Statistics from the model using all runs of data are shown. Distributions show the t-statistics of voxels from Non-Target (left) and Target (right) ROIs, defined as those that displayed significant positive stimulus-evoked changes relative to baseline (Non-Target) or in the contrast between Target and Non-Target conditions (Target) in the Standard data.

**Figure S2.**
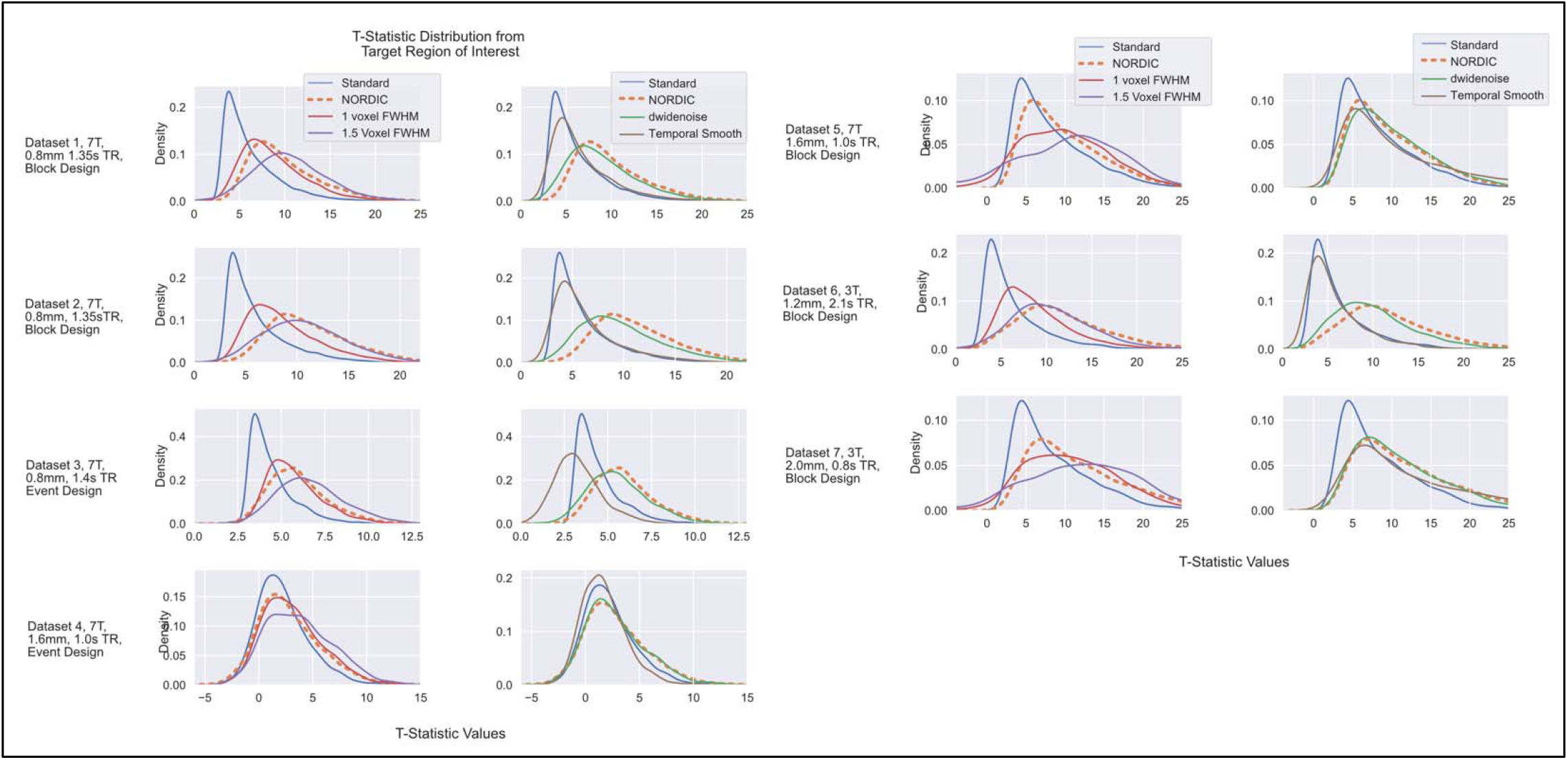
T-statistic values from Datasets 1 through 7 in the Target ROI, under different processing schemes within the Target ROI. Left column shows data from Standard, NORDIC and spatial smoothing with 1 and 1.5 voxel FWHM spatial smoothing. Right column compares the same Standard and NORDIC data against temporal smoothing and dwidenoise denoising. T-values were extracted from the Target ROI defined using the Standard data. The t-values obtained with NORDIC (Orange, dashed) processed data is comparable to the effects of an additional 1 or 1.5 voxels FWHM gaussian smoothing.

**Figure S3.**
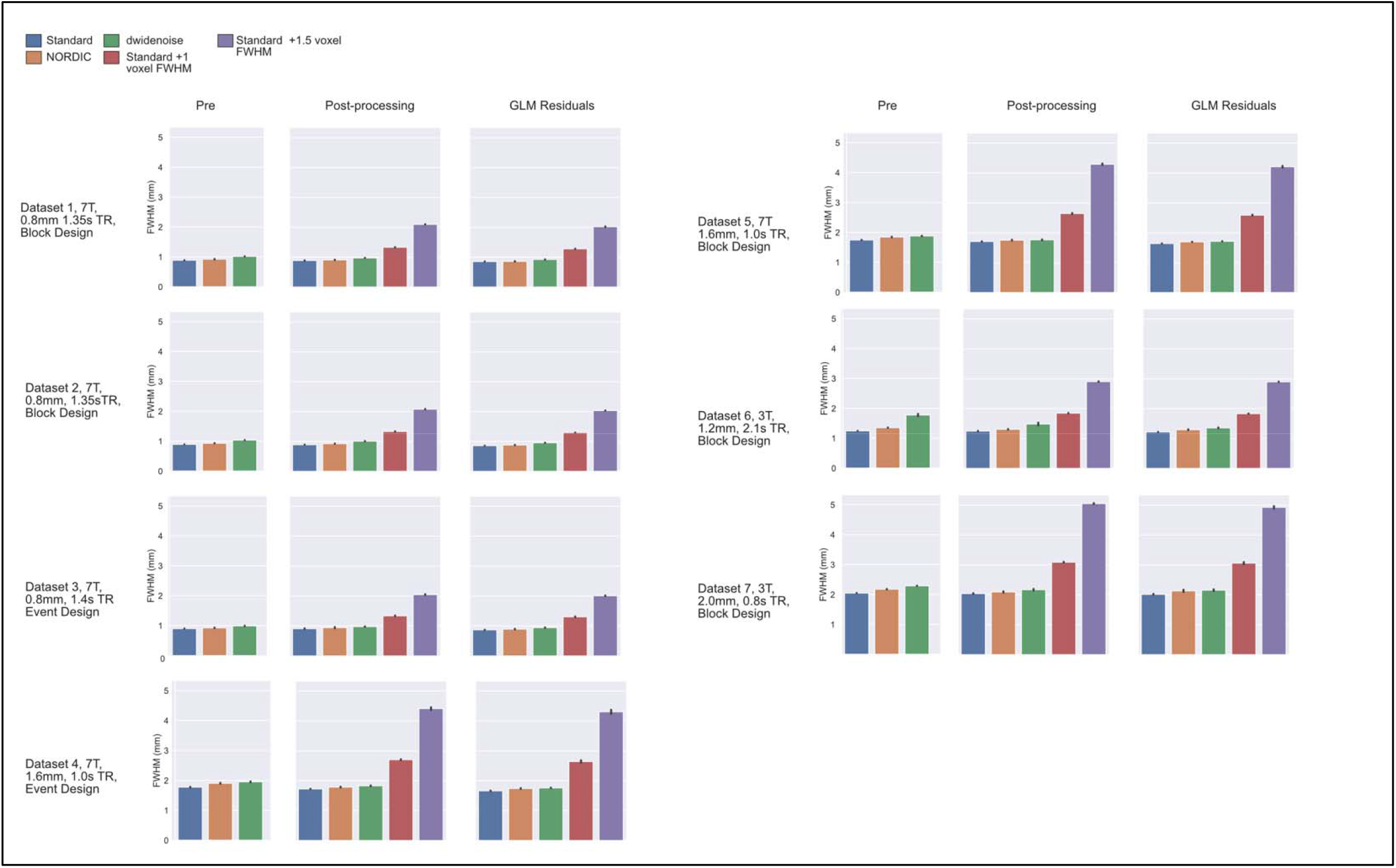
Global Smoothness Estimates for Datasets 1 – 7. Estimated spatial smoothness in mm (FWHM) at various processing stages for each method. Note that the image smoothness of the Standard, NORDIC, and dwidenoise data are substantially below the level of the additional 1 or 1.5 voxels of additional smoothing. These trends remain the same for the residuals (last columns) after the conventional GLM. Error bars indicate standard deviation over runs.

**Supplemental Figure S4.**
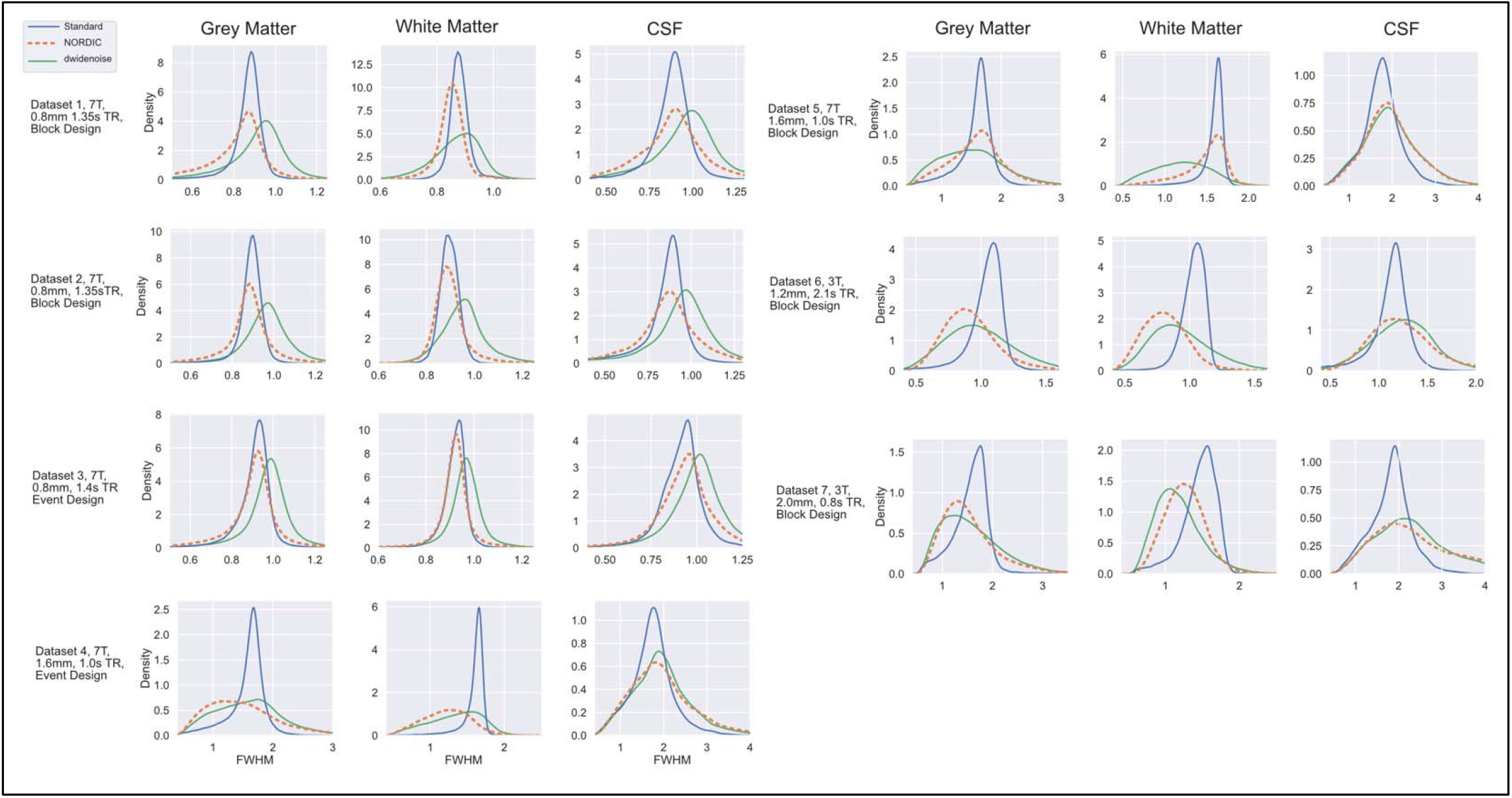
Local Smoothness from Datasets 1 – 7. These kernel density estimates show the distributions of the local spatial smoothness estimates in tissue classes derived from a T1-weighted anatomical image for Standard (blue), NORDIC (orange) and dwidenoise (green).

**Supplemental Figure S5.**
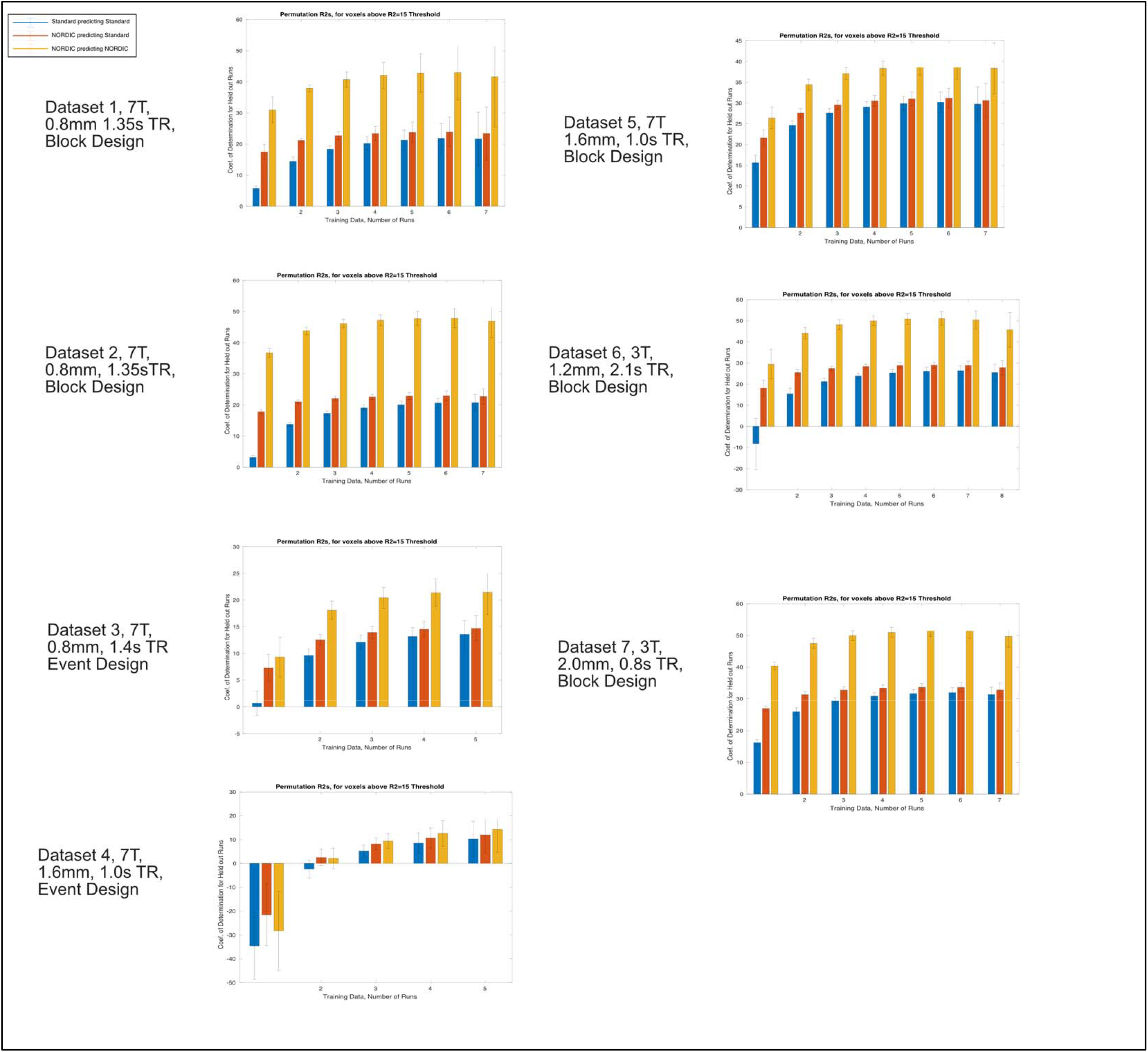
**Cross Validated R^2^ for Datasets 1 through 7** Leave-p-Out training was repeated for all vales of *p* greater than 1 and less than the number of runs. The number of runs included in the training vary across the X-axis, with bar height reflecting the R^2^ obtained. Including more data allows Standard models to approach, but not reach 2 to 3 runs of NORDIC data. Error bars indicate standard deviation.

**Supplemental Figure S6.**
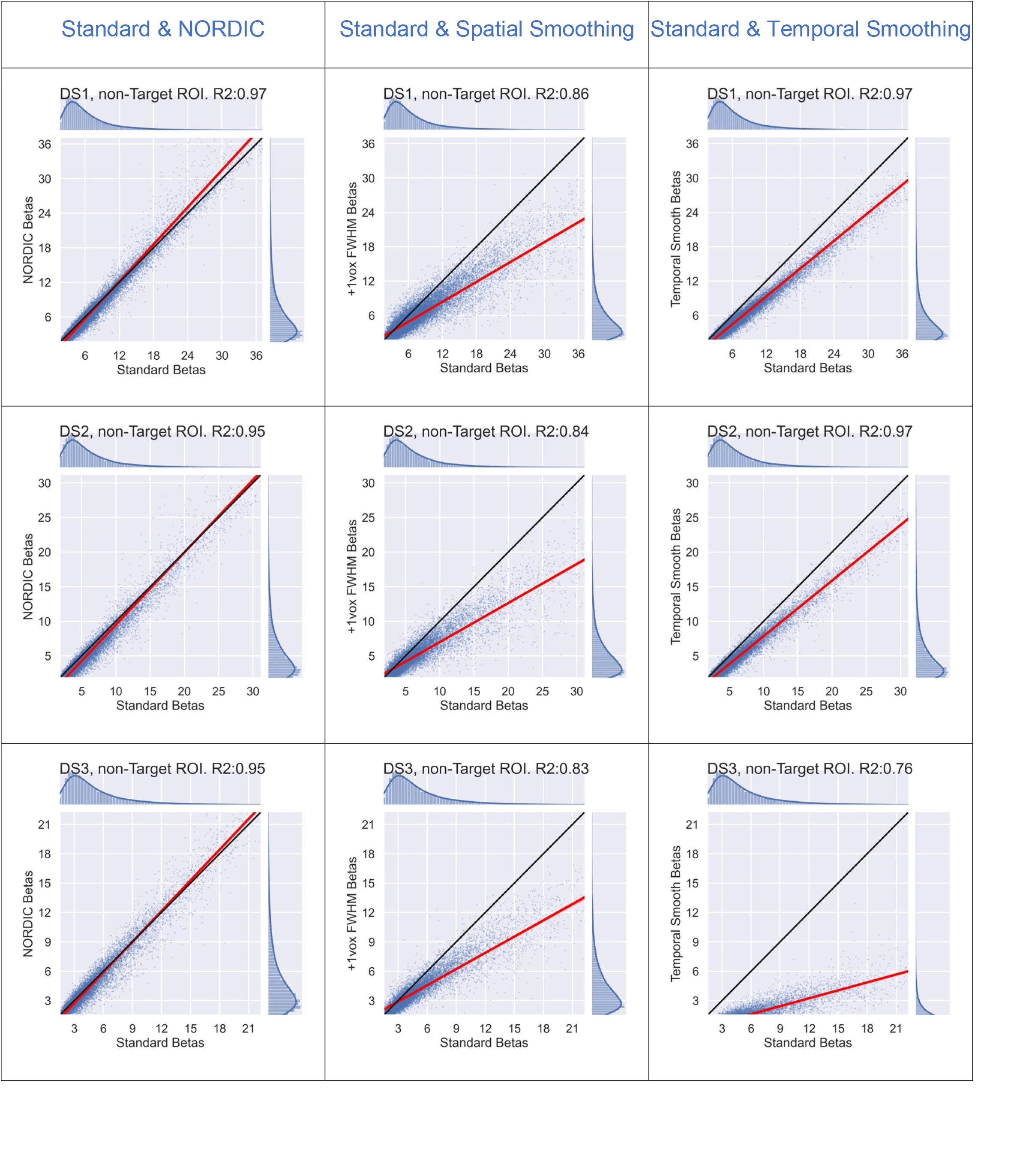
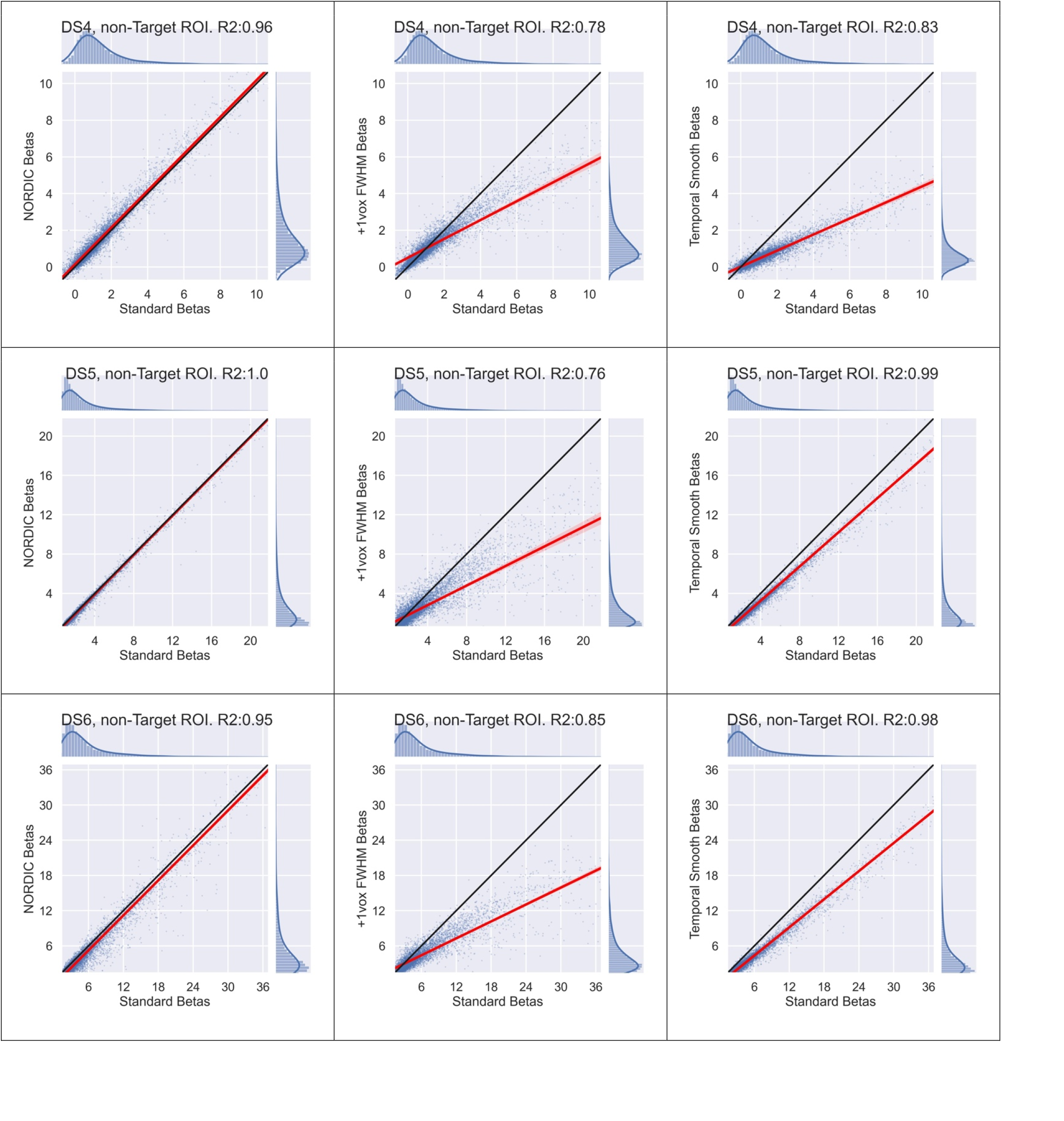
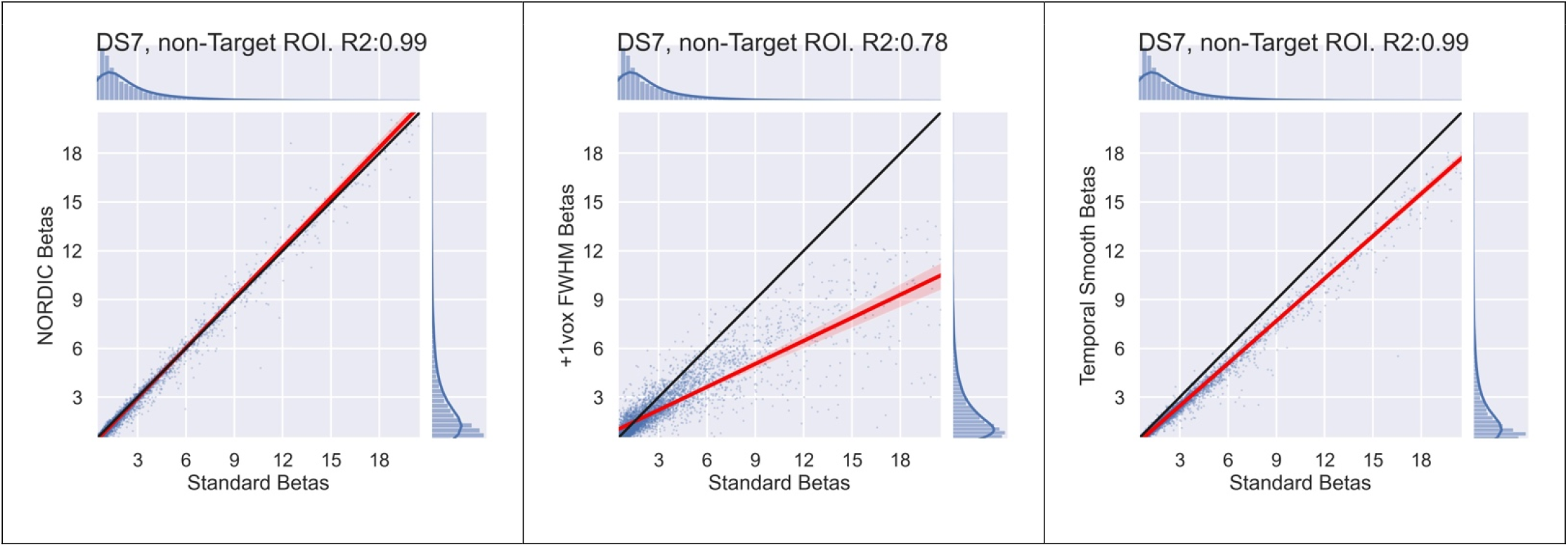
Scatter plots showing the relationship between the activation amplitude (i.e. beta, in percent signal change) within the large Non-Target ROI for the Standard data and NORDIC (1^st^ column), +1 Voxel spatial smoothing (2^nd^ column) or temporal smoothing (3^rd^ column). The black line is unity. The red line shows a regression line fit to the points. Distributions for each datatype are shown above and to the right to highlight that the vast majority of activation amplitudes are concentrated in the lower left-hand corner of the plot. The coefficient of determination, R^2^, is provided in each plot’s title.

### Local Perturbation Response analysis

Following the reviewers suggestion we implement the local perturbation response (LPR) method for evaluating non-linear reconstructions from Chan, C.C. and Haldar, J.P., 2021. We tested the LPR technique on in-vivo data, and also used it on a numerical simulation with random matrices. In LPR, a checker-board pattern of small amplitude is added to a single time-point, and for a measurement model, **Y**, the difference NORDIC (**Y**) - NORDIC (**Y**+LPR), in reconstruction is evaluated for the ability to **<u>recover</u>** the injected LPR signal and the effect of **<u>spreading</u>** of the injected LPR signal at other time-points.

In NORDIC the effect of the LPR can be tested on the hard threshold part for each patch by considering a model Y=X+N. If the model **X**, has a low-rank representation, then the LPR, which is simultaneously a low-rank and a sparse signal, is not necessarily aligned with the subspace containing **X**. Thus, intuitively only its projection onto this subspace can be recovered. It should be noted that if the additional LPR signal is expected to represent what is observed in fMRI data, its recovery may be tackled with robust PCA, designed for a low-rank + sparse model (Candès et al., 2011), but which has additional parameters as compared with hard thresholding. However, in our experience, we do not expect such vastly different patterns to be present for a single time-frame and vanish subsequently.

For the numerical simulation for Y=X+LPR +N, LPR was selected as a 36×36 checkerboard with 6×6 squares, both X and N had dimensions 1296 x 100, to maintain a ratio of 11:1. The entries of both X and N were i.i.d. and real valued distributed with variance 1.3, and 1 respectively, and 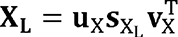 was the *R*-dimensional low-rank representation of 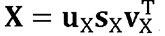, such that for n≤R the n singular-value 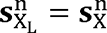 and for n>R, 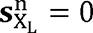. For the simulation, both the case of low-rank and full-rank model were evaluated with both separated and overlapping spectrum of singular values for the model and the added noise. As a quantitative metric for assessing the combination of noise and signal in time-points not probed by the LPR, the ratio 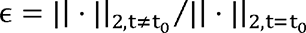 was used. The four cases of LPR recovery and spreading are shown in figure S7, along with their E value. For these cases, when X is low-rank the residual from LPR is more noticeable than when X is full-rank.

**Figure S7.**
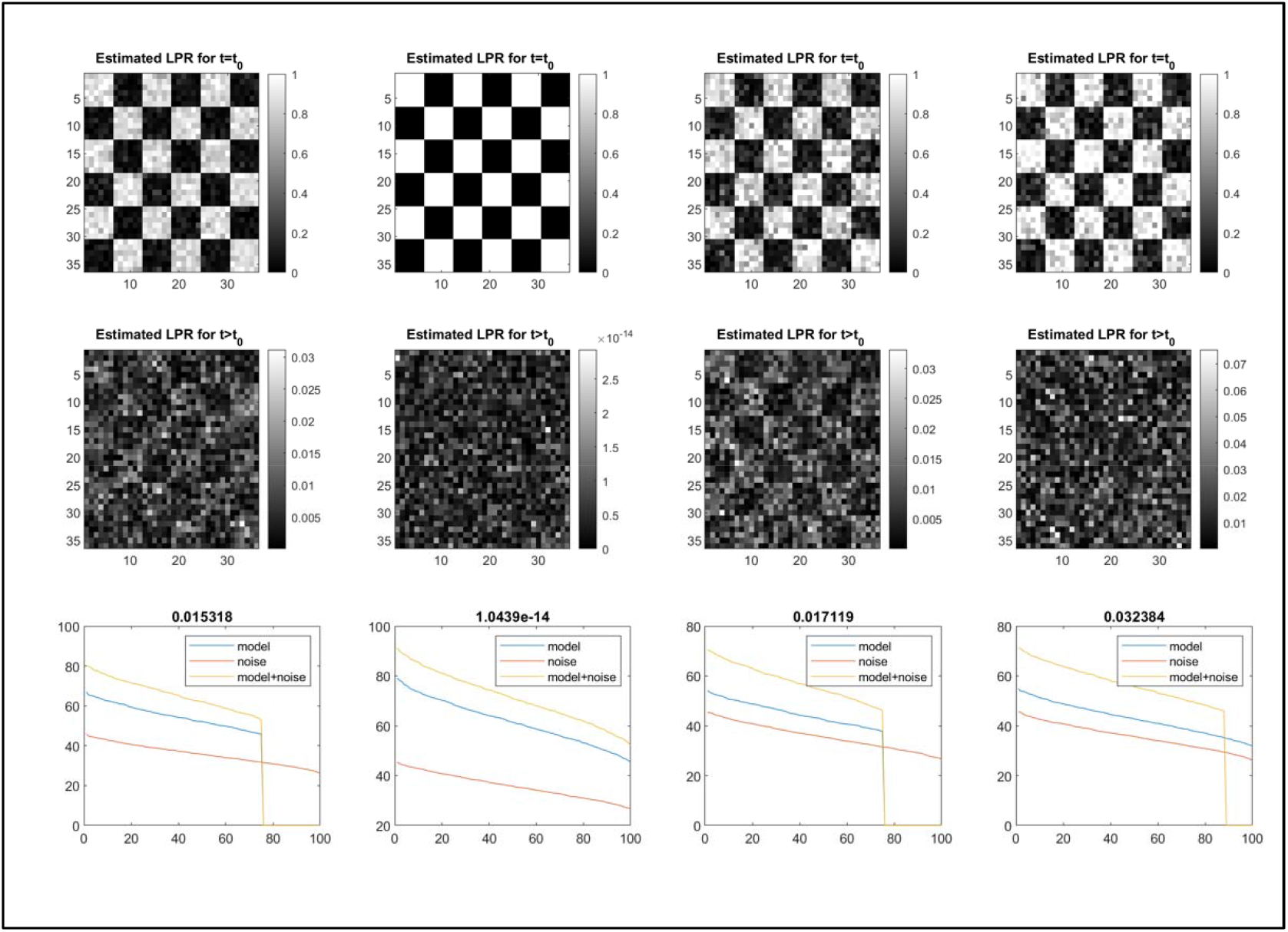
Numerical simulation. Four difference cases for utilizing an LPR (max(LPR)=σ)) are illustrated, for two models (low rank and full rank) and for two different noise-levels (overlapping and separated spectra of singular values).

We next added the LPR, at varying magnitudes relative to the measured thermal noise level, onto in vivo data, and performed NORDIC denoising. We then subtracted the original NORDIC data from the LPR+NORDIC data to examine to what extent the injected signal could be recovered following NORDIC.

When we consider all LPR intensities, we observe that the checkerboard LPR can be recovered, to some extent, even when it was of very low intensity relative to the thermal noise level of the data (Figure S8). At higher signal levels, recovery performance is increased and the checkerboard is clear.

**Figure S8.**
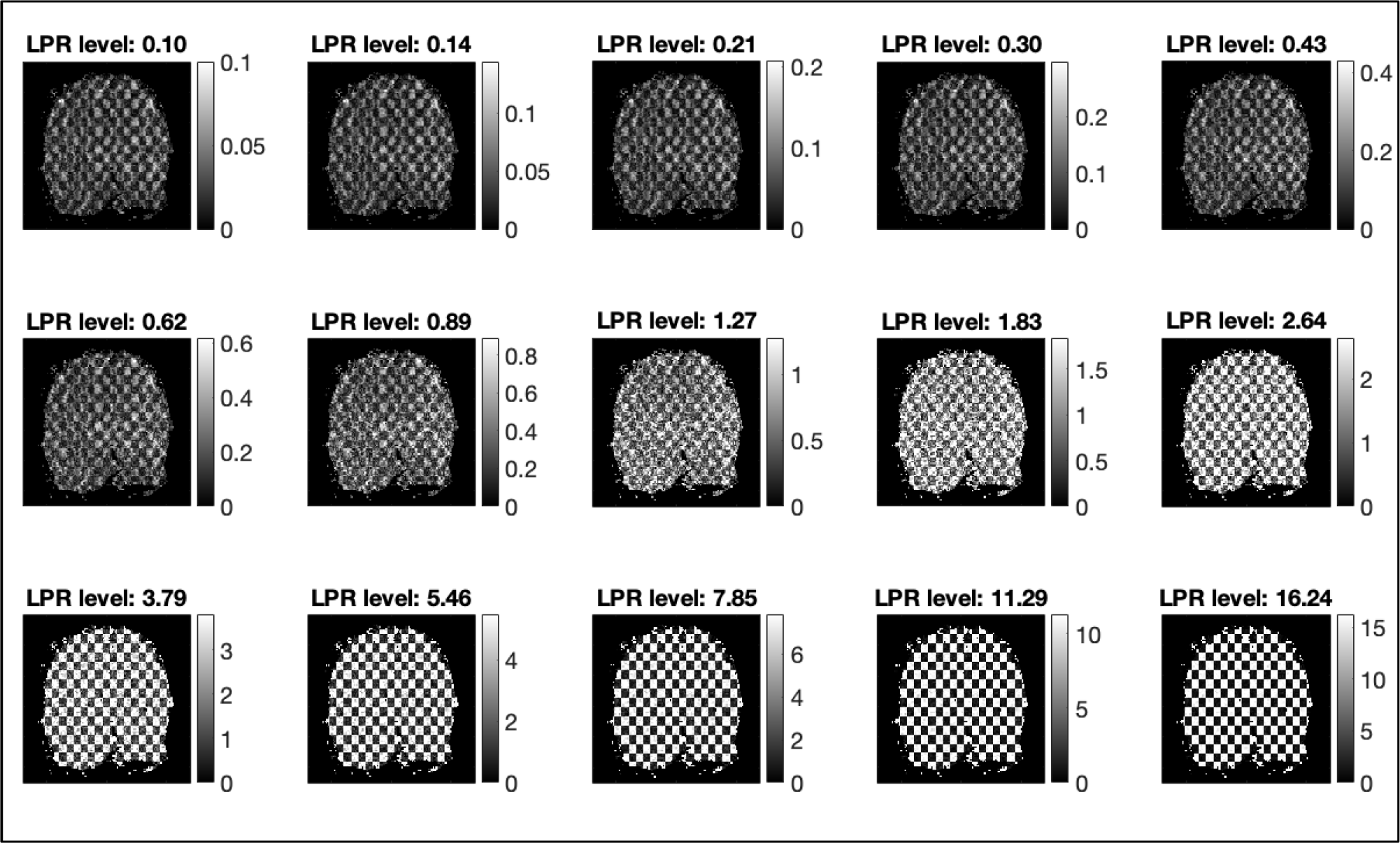
In vivo, recovery of injected LPR for different SNR levels. We observe that the LPR (a checkerboard) can be recovered even when the original LPR was very low in magnitude.

For a neighboring timepoint (Figure S9), we find that there is very limited artifactual signal from the injected LPR, with the highest relative energy at the lower LPR magnitudes. While there is some artifact just visible, the level of this artifact is order of magnitude lower than the original thermal noise or the fMRI signal fluctuations of interest.

**Figure S9.**
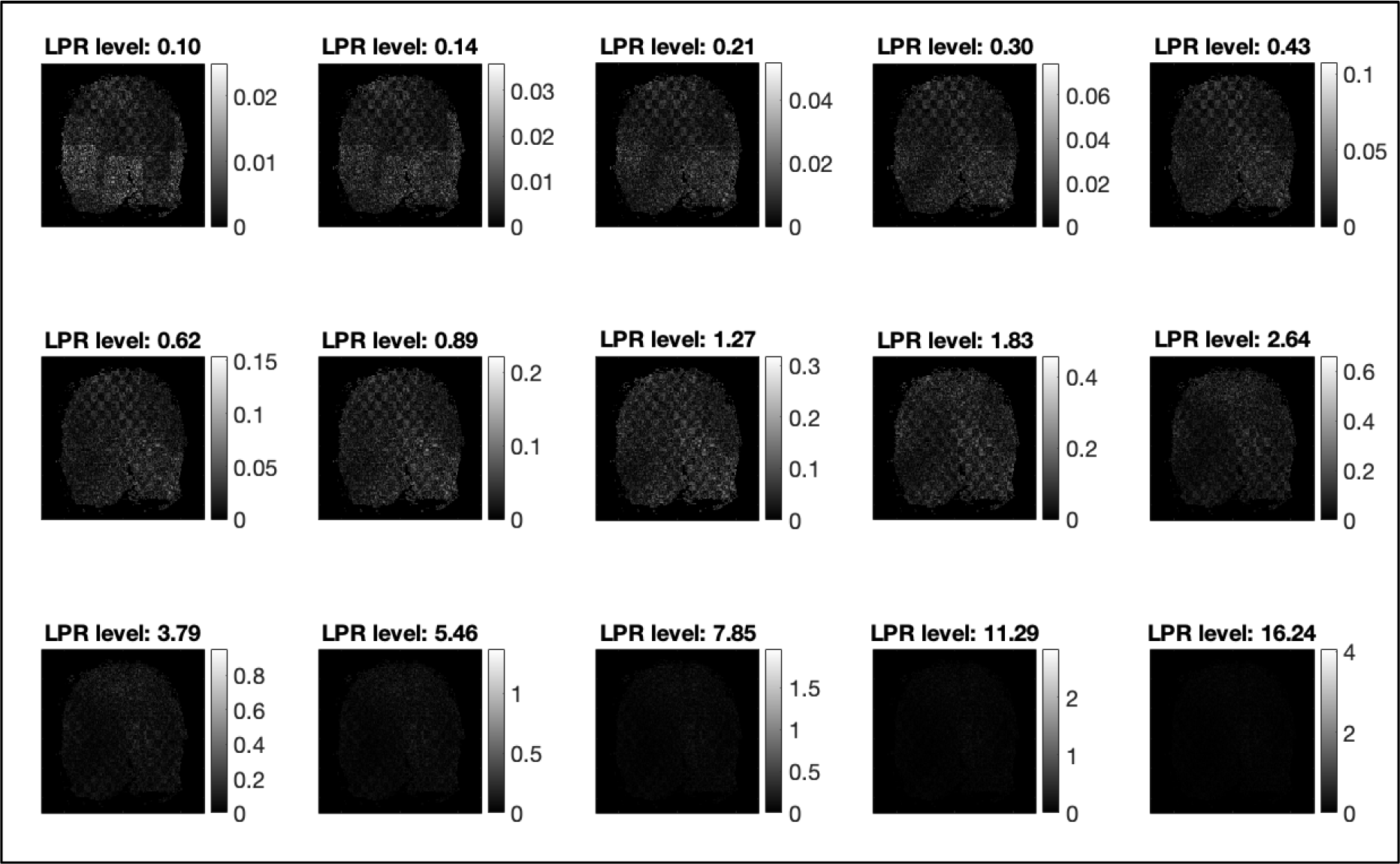
In vivo, comparison of neighboring timepoint after NORDIC in data with and without the LPR. The artifactual signal is at a very low intensity relative to the original injected LPR and primarily present at lower original LPR intensities (color map limits set to 25% of original LPR magnitude).

To examine the effect of this artifact, we can examine voxel time courses. Figure S10 shows the time course of 3 voxels for the original data (NO NORDIC), NORDIC and then NORDIC with 3 different LPR magnitudes. The largest effect is the suppression of thermal noise visible as the differences between the dashed black lines and the others. The effect of the spreading artifact would show up as differences between the blue lines and the 3 LPR levels – and is effectively invisible.

**Figure S10.** The minimal impact of the LPR artifact on voxel time courses. The black line shows the original data prior to NORDIC. Additional lines show the NORDIC data without the LPR (blue), and the NORDIC data with the injected LPR at various levels. While the artifact is measurable (Figures S11-S14) here we see that its effect is not meaningful. The time courses following NORDIC with and without the LPR are nearly indistinguishable.

To quantify this effect over all voxels and LPR magnitudes, we can consider the relationship between the magnitude of artifactual signal fluctuations and the magnitude of the signal fluctuations (Figure S11) following NORDIC (i.e. the temporal standard deviation). On average, this reaches a maximum of 0. Note that this means not that the artifact is causing 2% signal change, but rather that that the artifact is only 2% of the intrinsic fluctuations and thus causes negligible signal changes (as visible in the voxel time courses, Figure S10).

**Figure S11.**
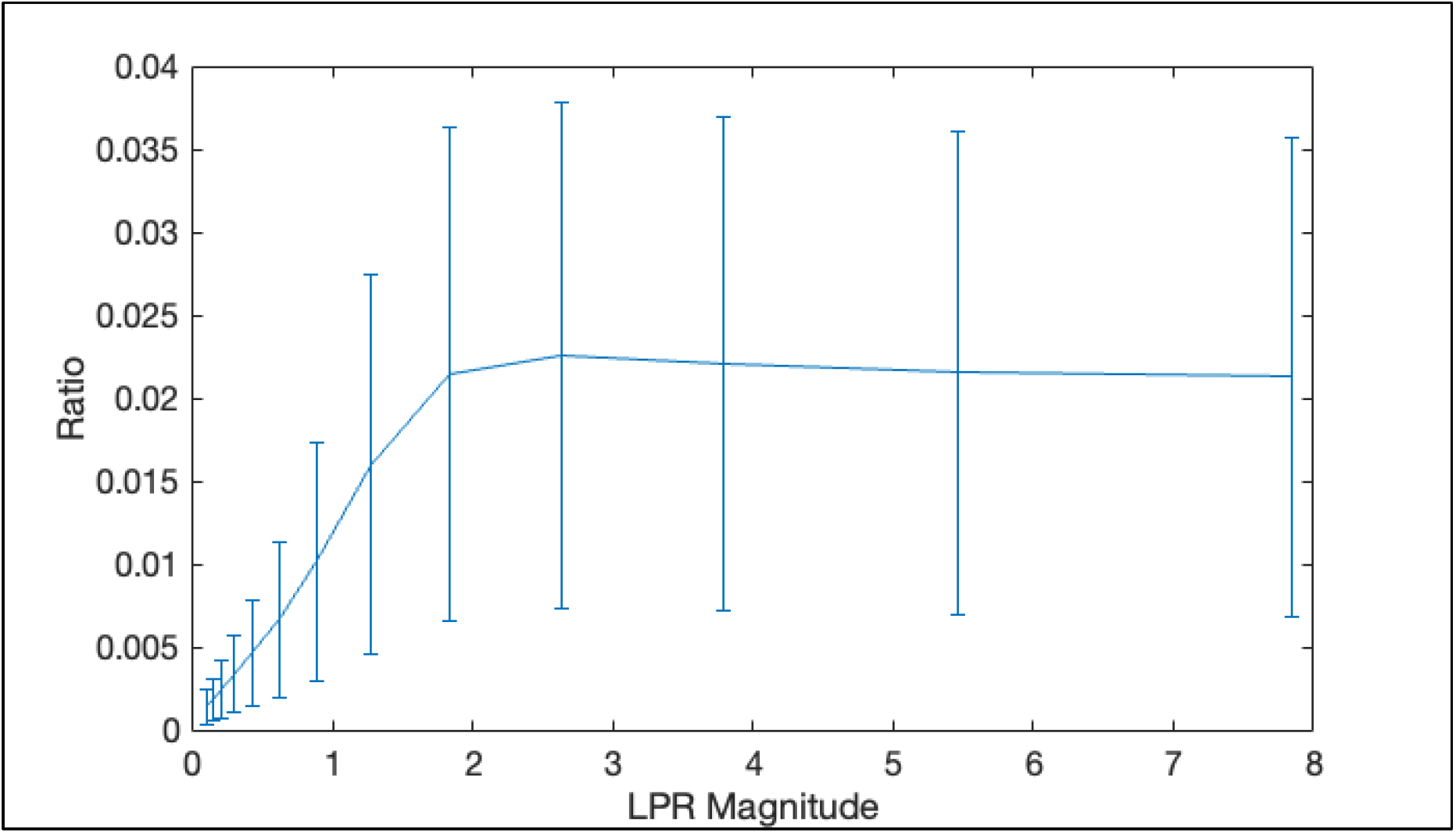
The magnitude of the artifactual fluctuations relative to intrinsic timeseries fluctuations. While Figures S12 and S13 showed the magnitude relative to the original LPR injection, here we are showing the magnitude of this artifact relative to the fluctuations in the denoised timeseries. While the artifact is visible (when data with and without the LPR are directly contrasted) its impact is minimal.

We can also summarize this as to the relationship of the recovered LPR and artifact signal to the original LPR signal. With an LPR for different thermal noise levels, the amount of energy recovered is close to the probed signal, and the residual is less than 1/10 of the probed signal, the plots of the simulation are shown in figure S12, with the in vivo signal shown in Figure S13.

**Figure S12.**
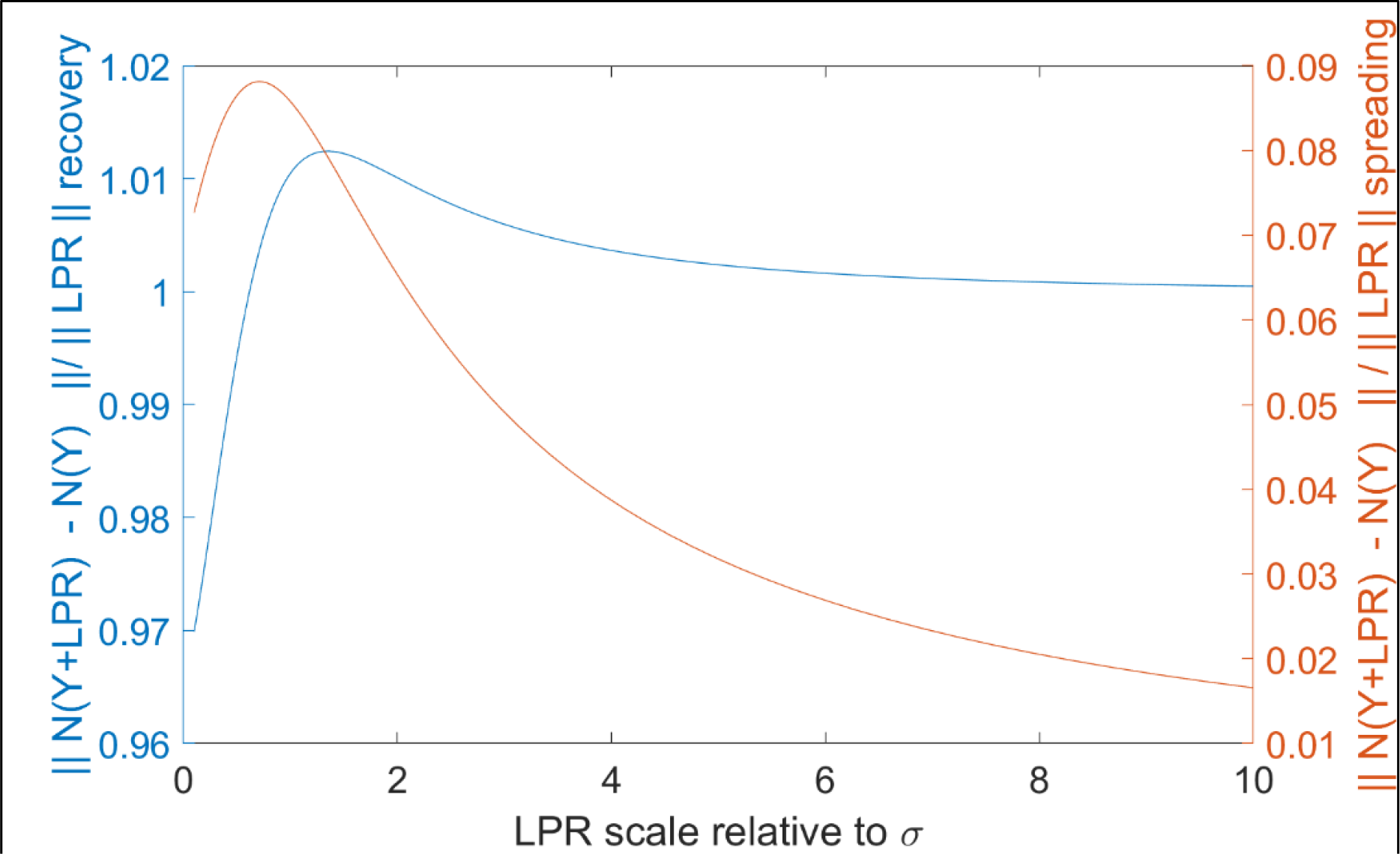
Plot of residual signal to LPR for different noise-regimes used in the simulation, showing recovery of LPR (blue) and LPR “artifact” at adjacent timepoints (orange).

**Figure S13.**
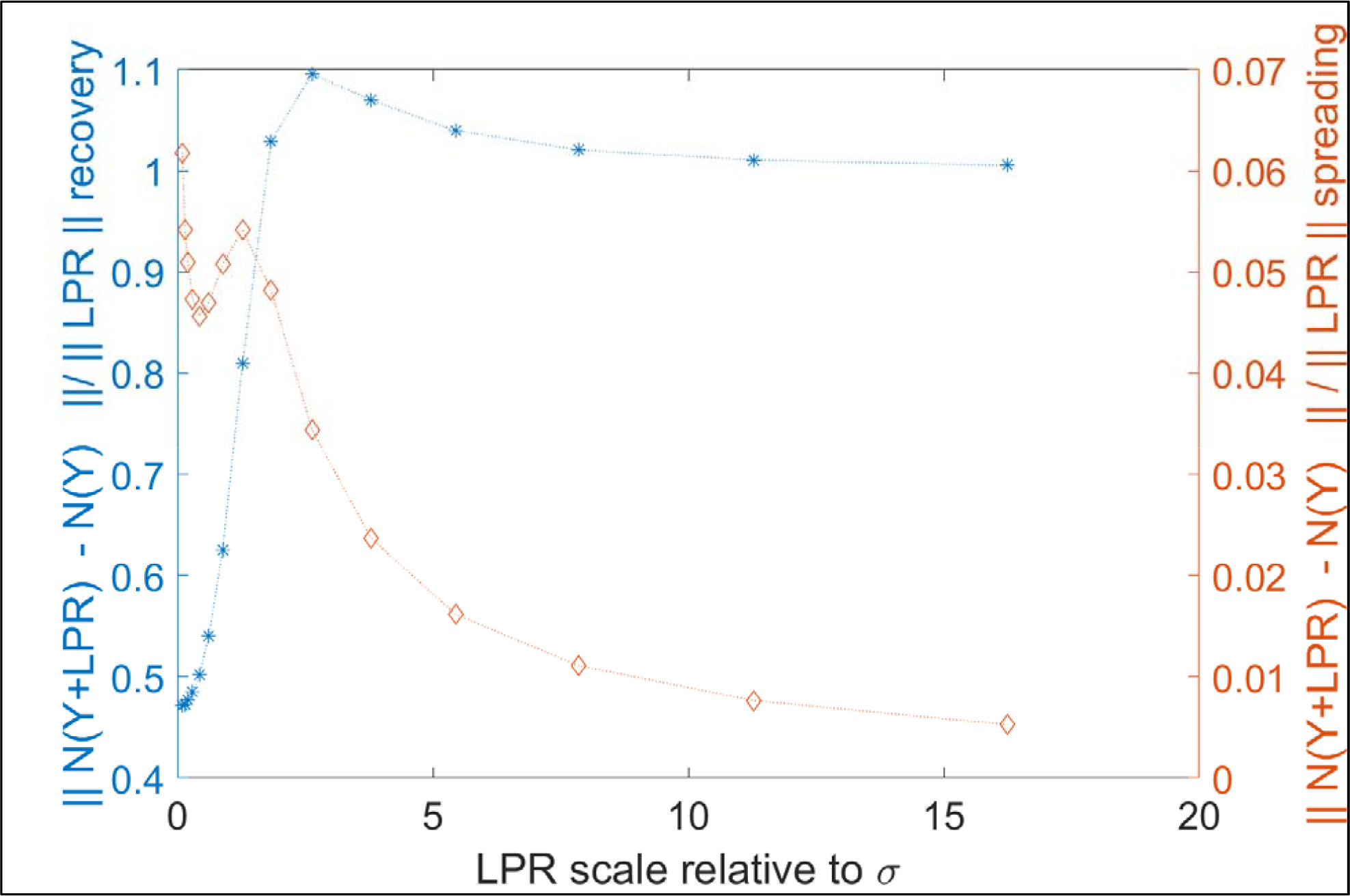
In vivo version of S8 showing recovery of LPR (blue) and LPR “artifact” at adjacent timepoints (orange).

From the numerical simulation the effect on the SVD of (X+LPR) vs the SVD of X for an LPR with a lower amplitude than the components in X is that the r first eigenvectors of both are almost the same, such that LPR is expressed into these basis functions, and then an r+1’th basis function is mostly identified with the remaining parts since it will be a “component”. This last basis function may or may not be recoverable, depending on the amplitude of the probed LPR. When recoverable, the representation of the LPR is the combination of the projection of the LPR onto the subspace spanned by X and any added basis function. The temporal sparse signal is likewise not described in a single explicit eigenvector but in the combination of eigenvectors. When the LPR is large, it is a large “peak” with noisy ripples for the primary eigenvector. When the LPR is low, the peak for representing a sparse signal is only achievable through the combination of eigenvectors.

A component not obtainable as being representable by 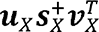 and which consistently is estimated as being (fully or partially) in 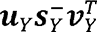, will persist in the final estimation of 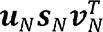. Such a residual component will be of less magnitude than 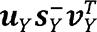 since for some patches it otherwise would be estimated as being in 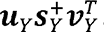. What is being discarded in NORDIC are those singular vector which are embedded in the distribution of the singular vectors of Gaussian noise. The associated eigenvectors are a low-rank representation of the observed full rank noise, and those eigenvectors are indiscriminately removed.

In combination the simulation and the in-vivo data shows that for probing date with a <u>sparse</u> signal at the noise level, using hard thresholding on the singular values for noise removal, a residual perturbation in the denoised signal at less than 1/10 the amplitude is observable, which reflects both that not all noise is removed, and that the model in NORDIC was chosen to recover low-rank signals.

### What do we mean by removing components of the timeseries which cannot be distinguished from Gaussian distributed noise?

For the SVD in NORDIC, the decomposition of the acquired signal may be written as

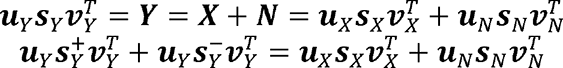

Where 𝑢 and 𝑣 are matrices with eigenvectors and where 𝑠_𝑋_ and 𝑠_𝑁_ are diagonal matrices with the singular values for the signal and noise respectively, and 𝑠_𝑁_ may be of full rank. The decomposition 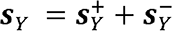 is such that 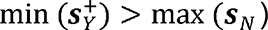 and 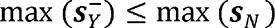. The estimated noise with hard thresholding in NORDIC is 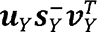, which is an approximation of the noise 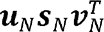, such that all the singular values in 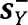 is less than the largest one in s_N_. It may be worth noting that the hard thresholding in NORDIC is lower than the optimal hard thresholding (Gavish and Donoho, 2014) or a low-rank signal. Likewise it may be informative to note that SVD is an orthonormal basis decomposition, where the observed signal (a row in ***Y***) is typically represented

by the combination of all eigenvectors in the decomposition of ***Y***, unless the decomposition happens to create an eigenvector that exactly matches such an observed signal. By extension the estimated eigenvectors for ***X*** will be impacted by the noise observed in ***Y*** and affecting the eigenvectors in the decomposition to most compactly model ***X***. In NORDIC, the basis functions which have an importance (i.e. corresponding singular value) less than what is observable from Gaussian noise is discarded.

**Supplemental Table S1.**
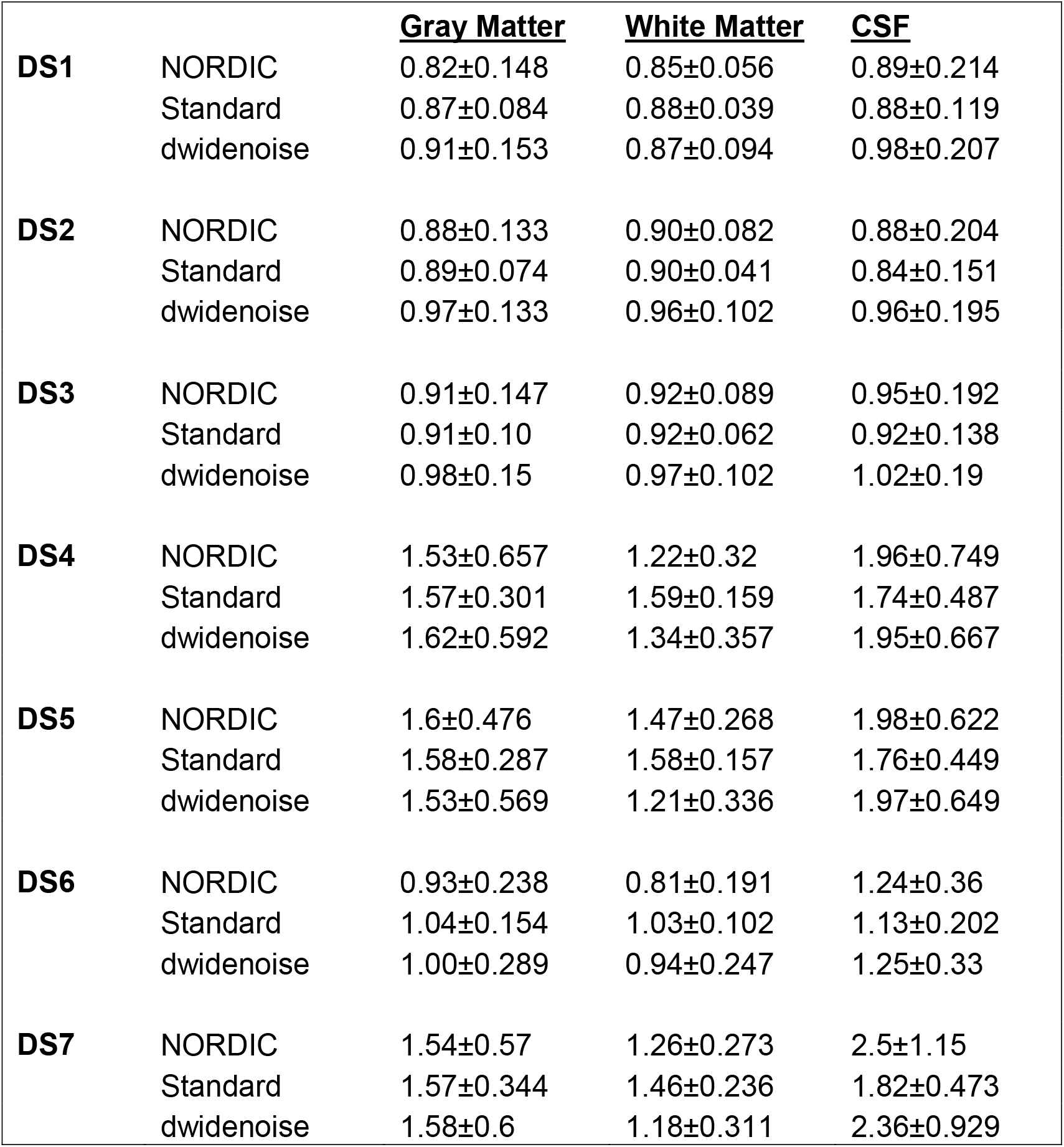
Mean and Standard Deviation of Local Smoothness Estimates in mm FWHM for Datasets 1 through 7.

**Supplemental Table 2.** The average Pearson correlations between the motion correction parameter estimates, with standard deviation over independent runs.

|  | **NORDIC, dwidenoise** | **Standard, dwidenoise** | **NORDIC, Standard** |
| --- | --- | --- | --- |
| **DS1** | 0.999±0.0008 | 0.999±0.0014 | 0.999±0.0012 |
| **DS2** | 0.997±0.003 | 0.992±0.0058 | 0.993±0.0054 |
| **DS3** | 0.998±0.0031 | 0.998±0.0024 | 0.997±0.0044 |
| **DS4** | 1±0.0001 | 1±0.0003 | 1±0.0003 |
| **DS5** | 1±0.0001 | 1±0.0002 | 1±0.0002 |
| **DS6** | 0.999±0.0018 | 0.993±0.0098 | 0.989±0.0173 |
| **DS7** | 1±0.0002 | 0.999±0.001 | 0.999±0.001 |

**Supplemental Table 3.** A comparison of the variability of FIR estimates within the target ROI for each dataset. This was calculated as the average (over voxels within the ROI mask) sum (over the time axis of the FIR) of the voxel-wise standard deviation over runs of the FIR response curves for the main task in each dataset (e.g. The center condition for DS1).

|  | **Standard Variability** | **NORDIC Variability** | **% Reduction** |
| --- | --- | --- | --- |
| **DS1** | 122.36 | 71.96 | 41.2 |
| **DS2** | 123.56 | 63.60 | 48.5 |
| **DS3** | 48.39 | 30.15 | 37.7 |
| **DS4** | 43.58 | 30.21 | 30.7 |
| **DS5** | 65.60 | 54.68 | 16.6 |
| **DS6** | 102.41 | 59.67 | 41.7 |
| **DS7** | 103.55 | 83.07 | 19.8 |

## References

1. Ades-Aron, B., Lemberskiy, G., Veraart, J., Golfinos, J., Fieremans, E., Novikov, D.S., Shepherd, T., 2021. Improved Task-based Functional MRI Language Mapping in Patients with Brain Tumors through Marchenko-Pastur Principal Component Analysis Denoising. Radiology 365–373.

2. Adhikari, B.M., Jahanshad, N., Shukla, D., Turner, J., Grotegerd, D., Dannlowski, U., Kugel, H., Engelen, J., Dietsche, B., Krug, A., Kircher, T., Fieremans, E., Veraart, J., Novikov, D.S., Boedhoe, P.S.W., van der Werf, Y.D., van den Heuvel, O.A., Ipser, J., Uhlmann, A., Stein, D.J., Dickie, E., Voineskos, A.N., Malhotra, A.K., Pizzagalli, F., Calhoun, V.D., Waller, L., Veer, I.M., Walter, H., Buchanan, R.W., Glahn, D.C., Hong, L.E., Thompson, P.M., Kochunov, P., 2019. A resting state fMRI analysis pipeline for pooling inference across diverse cohorts: an ENIGMA rs-fMRI protocol. Brain Imaging Behav. 13, 1453–1467. https://doi.org/10.1007/s11682-018-9941-x

3. Ashburner, J., Friston, K.J., 2005. Unified segmentation. NeuroImage 26, 839–851. https://doi.org/10.1016/j.neuroimage.2005.02.018

4. Bianciardi, M., van Gelderen, P., Duyn, J.H., Fukunaga, M., de Zwart, J.A., 2009. Making the most of fMRI at 7 T by suppressing spontaneous signal fluctuations. NeuroImage 448–454.

5. Blazejewska, A.I., Fischl, B., Wald, L.L., Polimeni, J.R., 2019. Intracortical smoothing of small-voxel fMRI data can provide increased detection power without spatial resolution losses compared to conventional large-voxel fMRI data. NeuroImage 189, 601–614. https://doi.org/10.1016/j.neuroimage.2019.01.054

6. Candès, E.J., Li, X., Ma, Y., Wright, J., 2011. Robust principal component analysis? J. ACM 58, 11:1–11:37. https://doi.org/10.1145/1970392.1970395

7. Candès, E.J., Sing-Long, C.A., Trzasko, J.D., 2013. Unbiased Risk Estimates for Singular Value Thresholding and Spectral Estimators. IEEE Trans. Signal Process. 61, 4643–4657. https://doi.org/10.1109/TSP.2013.2270464

8. Chan, C.C., Haldar, J.P., 2021. Chan, C.C., Haldar, J.P., 2021. Local perturbation responses and checkerboard tests: Characterization tools for nonlinear MRI methods. Magn Reson Med 86, 1873–1887. Magn. Reson. Med. 1873–1887.

9. Chen, G., Padmala, S., Chen, Y., Taylor, P.A., Cox, R.W., Pessoa, L., 2021. To pool or not to pool: Can we ignore cross-trial variability in FMRI? NeuroImage 225, 117496. https://doi.org/10.1016/j.neuroimage.2020.117496

10. Chen, W., Zhu, X.H., Thulborn, K.R., Ugurbil, K., 1999. Retinotopic mapping of lateral geniculate nucleus in humans using functional magnetic resonance imaging. Proc. Natl. Acad. Sci. 2430– 2434.

11. Cordero-Grande, L., Christiaens, D., Hutter, J., Price, A.N., Hajnal, J.V., 2019. Complex diffusion-weighted image estimation via matrix recovery under general noise models. NeuroImage 200, 391–404. https://doi.org/10.1016/j.neuroimage.2019.06.039

12. Cox, R.W., 1996. AFNI: Software for Analysis and Visualization of Functional Magnetic Resonance Neuroimages. Comput. Biomed. Res. 29, 162–173. https://doi.org/10.1006/cbmr.1996.0014

13. Cox, R.W., Chen, G., Glen, D.R., Reynolds, R.C., Taylor, P.A., 2017. FMRI Clustering in AFNI: False-Positive Rates Redux. Brain Connect. 7, 152–171. https://doi.org/10.1089/brain.2016.0475

14. De Martino, F., Yacoub, E., Kemper, V., Moerel, M., Uludağ, K., De Weerd, P., Ugurbil, K., Goebel, R., Formisano, E., 2018. The impact of ultra-high field MRI on cognitive and computational neuroimaging. NeuroImage, Neuroimaging with Ultra-high Field MRI: Present and Future 168, 366–382. https://doi.org/10.1016/j.neuroimage.2017.03.060

15. Dowdle, L.T., Ghose, G., Chen, C.C.C., Ugurbil, K., Yacoub, E., Vizioli, L., 2021a. Statistical power or more precise insights into neuro-temporal dynamics? Assessing the benefits of rapid temporal sampling in fMRI. Prog. Neurobiol., How high spatiotemporal resolution fMRI can advance neuroscience 207, 102171. https://doi.org/10.1016/j.pneurobio.2021.102171

16. Dowdle, L.T., Ghose, G., Ugurbil, K., Yacoub, E., Vizioli, L., 2021b. Clarifying the role of higher-level cortices in resolving perceptual ambiguity using ultra high field fMRI. NeuroImage 227, 117654. https://doi.org/10.1016/j.neuroimage.2020.117654

17. Dumoulin, S.O., Fracasso, A., van der Zwaag, W., Siero, J.C.W., Petridou, N., 2018. Ultra-high field MRI: Advancing systems neuroscience towards mesoscopic human brain function. NeuroImage, Neuroimaging with Ultra-high Field MRI: Present and Future 168, 345–357. https://doi.org/10.1016/j.neuroimage.2017.01.028

18. Duong, T.Q., Kim, D.S., Ugurbil, K., Kim, S.-G., 2001. Localized cerebral blood flow response at submillimeter columnar resolution. Proc. Natl. Acad. Sci. 98, 10904–10909.

19. DuPre, E., Salo, T., Ahmed, Z., Bandettini, P.A., Bottenhorn, K.L., Caballero-Gaudes, C., Dowdle, L.T., Gonzalez-Castillo, J., Heunis, S., Kundu, P., Laird, A.R., Markello, R., Markiewicz, C.J., Moia, S., Staden, I., Teves, J.B., Uruñuela, E., Vaziri-Pashkam, M., Whitaker, K., Handwerker, D.A., 2021. TE-dependent analysis of multi-echo fMRI with *tedana*. J. Open Source Softw. 6, 3669. https://doi.org/10.21105/joss.03669

20. Esteban, O., Markiewicz, C.J., Blair, R.W., Moodie, C.A., Isik, A.I., Erramuzpe, A., Kent, J.D., Goncalves, M., DuPre, E., Snyder, M., Oya, H., Ghosh, S.S., Wright, J., Durnez, J., Poldrack, R.A., Gorgolewski, K.J., 2019. fMRIPrep: a robust preprocessing pipeline for functional MRI. Nat. Methods 16, 111–116. https://doi.org/10.1038/s41592-018-0235-4

21. Finn, E.S., Huber, L., Bandettini, P.A., 2021. Higher and deeper: Bringing layer fMRI to association cortex. Prog. Neurobiol., How high spatiotemporal resolution fMRI can advance neuroscience 207, 101930. https://doi.org/10.1016/j.pneurobio.2020.101930

22. Friedman, L., Glover, G.H., Krenz, D., Magnotta, V., 2006. Reducing inter-scanner variability of activation in a multicenter fMRI study: Role of smoothness equalization. NeuroImage 32, 1656–1668. https://doi.org/10.1016/j.neuroimage.2006.03.062

23. Friedman, L., Stern, H., Brown, G.G., Mathalon, D.H., Turner, J., Glover, G.H., Gollub, R.L., Lauriello, J., Lim, K.O., Cannon, T., Greve, D.N., Bockholt, H.J., Belger, A., Mueller, B., Doty, M.J., He, J., Wells, W., Smyth, P., Pieper, S., Kim, S., Kubicki, M., Vangel, M., Potkin, S.G., 2008. Test– retest and between-site reliability in a multicenter fMRI study. Hum. Brain Mapp. 29, 958–972. https://doi.org/10.1002/hbm.20440

24. Gavish, M., Donoho, D.L., 2014. The Optimal Hard Threshold for Singular Values is $4/\sqrt 3$. IEEE Trans. Inf. Theory 60, 5040–5053. https://doi.org/10.1109/TIT.2014.2323359

25. Glover, G.H., Li, T.-Q., Ress, D., 2000. Image-based method for retrospective correction of physiological motion effects in fMRI: RETROICOR. Magn. Reson. Med. 44, 162–167. https://doi.org/10.1002/1522-2594(200007)44:1<162::AID-MRM23>3.0.CO;2-E

26. Gonzalez-Castillo, J., Panwar, P., Buchanan, L.C., Caballero-Gaudes, C., Handwerker, D.A., Jangraw, D.C., Zachariou, V., Inati, S., Roopchansingh, V., Derbyshire, J.A., Bandettini, P.A., 2016. Evaluation of multi-echo ICA denoising for task based fMRI studies: Block designs, rapid event-related designs, and cardiac-gated fMRI. Neuroimage 141, 452–468. https://doi.org/10.1016/j.neuroimage.2016.07.049

27. Gonzalez-Castillo, J., Saad, Z.S., Handwerker, D.A., Inati, S.J., Brenowitz, N., Bandettini, P.A., 2012. Whole-brain, time-locked activation with simple tasks revealed using massive averaging and model-free analysis. Proc. Natl. Acad. Sci. U. S. A. 109, 5487–5492. https://doi.org/10.1073/pnas.1121049109

28. Haldar, J.P., Liang, Z.-P., 2011. Low-rank approximations for dynamic imaging, in: 2011 IEEE International Symposium on Biomedical Imaging: From Nano to Macro. Presented at the 2011 IEEE International Symposium on Biomedical Imaging: From Nano to Macro, pp. 1052–1055. https://doi.org/10.1109/ISBI.2011.5872582

29. Handwerker, D.A., Ollinger, J.M., D’Esposito, M., 2004. Variation of BOLD hemodynamic responses across subjects and brain regions and their effects on statistical analyses. NeuroImage 21, 1639–1651. https://doi.org/10.1016/j.neuroimage.2003.11.029

30. Haxby, J.V., Connolly, A.C., Guntupalli, J.S., 2014. Decoding Neural Representational Spaces Using Multivariate Pattern Analysis. Annu. Rev. Neurosci. 37, 435–456. https://doi.org/10.1146/annurev-neuro-062012-170325

31. Hu, X., Kim, S.G., 1994. Reduction of signal fluctuation in functional MRI using navigator echoes. Magn. Reson. Med. 31, 495–503.

32. Huber, L., Finn, E.S., Handwerker, D.A., Bönstrup, M., Glen, D.R., Kashyap, S., Ivanov, D., Petridou, N., Marrett, S., Goense, J., Poser, B.A., Bandettini, P.A., 2020. Sub-millimeter fMRI reveals multiple topographical digit representations that form action maps in human motor cortex. NeuroImage 208, 116463. https://doi.org/10.1016/j.neuroimage.2019.116463

33. Huber, L., Ivanov, D., Handwerker, D.A., Marrett, S., Guidi, M., Uludağ, K., Bandettini, P.A., Poser, B.A., 2018. Techniques for blood volume fMRI with VASO: From low-resolution mapping towards sub-millimeter layer-dependent applications. NeuroImage, Pushing the spatio-temporal limits of MRI and fMRI 164, 131–143. https://doi.org/10.1016/j.neuroimage.2016.11.039

34. Huber, L., Poser, B.A., Bandettini, P.A., Arora, K., Wagstyl, K., Cho, S., Shinho Cho, Shinho Cho, Goense, J., Nothnagel, N., Morgan, A.T., van den Hurk, J., Müller, A.K., Reynolds, R.C., Glen, D.R., Goebel, R., Gulban, O.F., 2021. LayNii: A software suite for layer-fMRI. NeuroImage 237, 118091–118091. https://doi.org/10.1016/j.neuroimage.2021.118091

35. Kay, K., Rokem, A., Winawer, J., Dougherty, R., Wandell, B., 2013. GLMdenoise: a fast, automated technique for denoising task-based fMRI data. Front. Neurosci. 7. https://doi.org/10.3389/fnins.2013.00247

36. Kay, K.N., Naselaris, T., Prenger, R.J., Gallant, J.L., 2008. Identifying natural images from human brain activity. Nature 325–355.

37. Kim, S.G., Hendrich, K., Hu, X., Merkle, H., Ugurbil, K., 1994. Potential pitfalls of functional MRI using conventional gradient-recalled echo techniques. NMR Biomed. 69–74.

38. Kundu, P., Voon, V., Balchandani, P., Lombardo, M.V., Poser, B.A., Bandettini, P., 2017. Multi-Echo fMRI: A Review of Applications in fMRI Denoising and Analysis of BOLD Signals. Neuroimage. https://doi.org/10.1016/j.neuroimage.2017.03.033

39. Lawrence, S.J.D., Formisano, E., Muckli, L., de Lange, F.P., 2019. Laminar fMRI: Applications for cognitive neuroscience. NeuroImage 197, 785–791. https://doi.org/10.1016/j.neuroimage.2017.07.004

40. Lund, T.E., Madsen, K.H., Sidaros, K., Luo, W.L., Nichols, T.E., 2006. Non-white noise in fMRI: does modelling have an impact? NeuroImage 54–66.

41. Marcus, D.S., Harms, M.P., Snyder, A.Z., Jenkinson, M., Wilson, J.A., Glasser, M.F., Barch, D.M., Archie, K.A., Burgess, G.C., Ramaratnam, M., Hodge, M., Horton, W., Herrick, R., Olsen, T., McKay, M., House, M., Hileman, M., Reid, E., Harwell, J., Coalson, T., Schindler, J., Elam, J.S., Curtiss, S.W., Van Essen, D.C., 2013. Human Connectome Project informatics: Quality control, database services, and data visualization. NeuroImage, Mapping the Connectome 80, 202–219. https://doi.org/10.1016/j.neuroimage.2013.05.077

42. Marques, J.P., Kober, T., Krueger, G., van der Zwaag, W., Van de Moortele, P.-F., Gruetter, R., 2010. MP2RAGE, a self bias-field corrected sequence for improved segmentation and T1-mapping at high field. NeuroImage 49, 1271–1281. https://doi.org/10.1016/j.neuroimage.2009.10.002

43. Meyer, N.K., Campeau, N.G., Black, D.F., Welker, K.M., Gunter, J.L., Yarach, U., Kang, D., In, M., Huston III, J., Shu, Y., Bernstein, M.A., Trzasko, J.D., 2020. Locally low-rank denoising of complex-valued EPI reconstructions preceding task fMRI analysis. Presented at the ISMRM, Virtual.

44. Moeller, S., Pisharady, P.K., Ramanna, S., Lenglet, C., Wu, X., Dowdle, L., Yacoub, E., Uğurbil, K., Akçakaya, M., 2021. NOise reduction with DIstribution Corrected (NORDIC) PCA in dMRI with complex-valued parameter-free locally low-rank processing. NeuroImage 226, 117539. https://doi.org/10.1016/j.neuroimage.2020.117539

45. Moeller, S., Yacoub, E., Olman, C.A., Auerbach, E., Strupp, J., Harel, N., Uğurbil, K., 2010. Multiband multislice GE-EPI at 7 tesla, with 16-fold acceleration using partial parallel imaging with application to high spatial and temporal whole-brain fMRI. Magn. Reson. Med. 63, 1144–1153. https://doi.org/10.1002/mrm.22361

46. Mugler, J.P., Brookeman, J.R., 1991. Rapid three-dimensional T1-weighted MR imaging with the MP-RAGE sequence. J. Magn. Reson. Imaging JMRI 1, 561–567. https://doi.org/10.1002/jmri.1880010509

47. Naselaris, T., Kay, K.N., Nishimoto, S., Gallant, J.L., 2011. Encoding and decoding in fMRI. NeuroImage 400–410.

48. Norris, D.G., Polimeni, J.R., 2019. Laminar (f)MRI: A short history and future prospects. NeuroImage 197, 643–649. https://doi.org/10.1016/j.neuroimage.2019.04.082

49. Olszowy, W., Aston, J., Rua, C., Williams, G.B., 2019. Accurate autocorrelation modeling substantially improves fMRI reliability. Nat. Commun. 10, 1–11. https://doi.org/10.1038/s41467-019-09230-w

50. Polimeni, J.R., Lewis, L.D., 2021. Imaging faster neural dynamics with fast fMRI: A need for updated models of the hemodynamic response. Prog. Neurobiol., How high spatiotemporal resolution fMRI can advance neuroscience 207, 102174. https://doi.org/10.1016/j.pneurobio.2021.102174

51. Polimeni, J.R., Renvall, V., Zaretskaya, N., Fischl, B., 2018. Analysis strategies for high-resolution UHF-fMRI data. NeuroImage, Neuroimaging with Ultra-high Field MRI: Present and Future 168, 296–320. https://doi.org/10.1016/j.neuroimage.2017.04.053

52. Polimeni, J.R., Uludağ, K., 2018. Neuroimaging with ultra-high field MRI: Present and future. NeuroImage, Neuroimaging with Ultra-high Field MRI: Present and Future 168, 1–6. https://doi.org/10.1016/j.neuroimage.2018.01.072

53. Pruessmann, K.P., Weiger, M., Scheidegger, M.B., Boesiger, P., 1999. SENSE: sensitivity encoding for fast MRI. Magn. Reson. Med. 42, 952–962.

54. Pruim, R.H.R., Mennes, M., Buitelaar, J.K., Beckmann, C.F., 2015. Evaluation of ICA-AROMA and alternative strategies for motion artifact removal in resting state fMRI. NeuroImage 112, 278–287. https://doi.org/10.1016/j.neuroimage.2015.02.063

55. Roberts, D.A., Detre, J.A., Bolinger, L., Insko, E.K., Leigh, J.S., 1994. Quantitative magnetic resonance imaging of human brain perfusion at 1.5 T using steady-state inversion of arterial water. Proc. Natl. Acad. Sci. 91, 33–37. https://doi.org/10.1073/pnas.91.1.33

56. Saad, Z.S., Glen, D.R., Chen, G., Beauchamp, M.S., Desai, R., Cox, R.W., 2009. A new method for improving functional-to-structural MRI alignment using local Pearson correlation. NeuroImage 44, 839–848. https://doi.org/10.1016/j.neuroimage.2008.09.037

57. Shmuel, A., Yacoub, E., Chaimow, D., Logothetis, N.K., Ugurbil, K., 2007. Spatio-temporal point-spread function of fMRI signal in human gray matter at 7 Tesla. NeuroImage 35, 539–552. https://doi.org/10.1016/j.neuroimage.2006.12.030

58. Shmueli, K., van Gelderen, P., de Zwart, J.A., Horovitz, S.G., Fukunaga, M., Jansma, J.M., Duyn, J.H., 2007. Low-frequency fluctuations in the cardiac rate as a source of variance in the resting-state fMRI BOLD signal. NeuroImage 306–320.

59. Smith, S.M., Beckmann, C.F., Andersson, J., Auerbach, E.J., Bijsterbosch, J., Douaud, G., Duff, E., Feinberg, D.A., Griffanti, L., Harms, M.P., Kelly, M., Laumann, T., Miller, K.L., Moeller, S., Petersen, S., Power, J., Salimi-Khorshidi, G., Snyder, A.Z., Vu, A.T., Woolrich, M.W., Xu, J., Yacoub, E., Uğurbil, K., Van Essen, D.C., Glasser, M.F., 2013. Resting-state fMRI in the Human Connectome Project. NeuroImage, Mapping the Connectome 80, 144–168. https://doi.org/10.1016/j.neuroimage.2013.05.039

60. Smith, S.M., Fox, P.T., Miller, K.L., Glahn, D.C., Fox, P.M., Mackay, C.E., Filippini, N., Watkins, K.E., Toro, R., Laird, A.R., Beckmann, C.F., 2009. Correspondence of the brain’s functional architecture during activation and rest. Proc. Natl. Acad. Sci. 13040–13045.

61. Stringer, E.A., Chen, L.M., Friedman, R.M., Gatenby, C., Gore, J.C., 2011. Stringer, E.A., Chen, L.M., Friedman, R.M., Gatenby, C., Gore, J.C., 2011. Differentiation of somatosensory cortices by high-resolution fMRI at 7 T. NeuroImage 54, 1012–1020.

62. Taylor, A.J., Kim, J.H., Ress, D., 2018. Characterization of the hemodynamic response function across the majority of human cerebral cortex. NeuroImage 173, 322–331. https://doi.org/10.1016/j.neuroimage.2018.02.061

63. Thomas, C.G., Harshman, R.A., Menon, R.S., 2002. Noise Reduction in BOLD-Based fMRI Using Component Analysis. NeuroImage 17, 1521–1537. https://doi.org/10.1006/nimg.2002.1200

64. Todd, N., Josephs, O., Zeidman, P., Flandin, G., Moeller, S., Weiskopf, N., 2017. Functional Sensitivity of 2D Simultaneous Multi-Slice Echo-Planar Imaging: Effects of Acceleration on g-factor and Physiological Noise. Front. Neurosci. 11. https://doi.org/10.3389/fnins.2017.00158

65. Tournier, J.-D., Smith, R., Raffelt, D., Tabbara, R., Dhollander, T., Pietsch, M., Christiaens, D., Jeurissen, B., Yeh, C.-H., Connelly, A., 2019. MRtrix3: A fast, flexible and open software framework for medical image processing and visualisation. NeuroImage 202, 116137. https://doi.org/10.1016/j.neuroimage.2019.116137

66. Triantafyllou, C., Hoge, R.D., Krueger, G., Wiggins, C.J., Potthast, A., Wiggins, G.C., Wald, L.L., 2005. Comparison of physiological noise at 1.5 T, 3 T and 7 T and optimization of fMRI acquisition parameters. NeuroImage 26, 243–250. https://doi.org/10.1016/j.neuroimage.2005.01.007

67. Triantafyllou, C., Hoge, R.D., Wald, L.L., 2006. Effect of spatial smoothing on physiological noise in high-resolution fMRI. NeuroImage 551–557.

68. Triantafyllou, C., Polimeni, J.R., Wald, L.L., 2011. Physiological noise and signal-to-noise ratio in fMRI with multi-channel array coils. Neuroimage 55, 597–606. https://doi.org/10.1016/j.neuroimage.2010.11.084

69. Uğurbil, K., 2018. Imaging at ultrahigh magnetic fields: History, challenges, and solutions. NeuroImage, Neuroimaging with Ultra-high Field MRI: Present and Future 168, 7–32. https://doi.org/10.1016/j.neuroimage.2017.07.007

70. Ugurbil, K., 2016. What is feasible with imaging human brain function and connectivity using functional magnetic resonance imaging. Philos. Trans. R. Soc. B Biol. Sci. 371, 20150361. https://doi.org/10.1098/rstb.2015.0361

71. Uğurbil, K., 2014. Magnetic Resonance Imaging at Ultrahigh Fields. IEEE Trans. Biomed. Eng. 61, 1364–1379. https://doi.org/10.1109/TBME.2014.2313619

72. Uğurbil, K., Xu, J., Auerbach, E.J., Moeller, S., Vu, A.T., Duarte-Carvajalino, J.M., Lenglet, C., Wu, X., Schmitter, S., Van de Moortele, P.F., Strupp, J., Sapiro, G., De Martino, F., Wang, D., Harel, N., Garwood, M., Chen, L., Feinberg, D.A., Smith, S.M., Miller, K.L., Sotiropoulos, S.N., Jbabdi, S., Andersson, J.L.R., Behrens, T.E.J., Glasser, M.F., Van Essen, D.C., Yacoub, E., 2013. Pushing spatial and temporal resolution for functional and diffusion MRI in the Human Connectome Project. NeuroImage, Mapping the Connectome 80, 80–104. https://doi.org/10.1016/j.neuroimage.2013.05.012

73. Veraart, J., Novikov, D.S., Christiaens, D., Ades-aron, B., Sijbers, J., Fieremans, E., 2016. Denoising of diffusion MRI using random matrix theory. NeuroImage 142, 394–406. https://doi.org/10.1016/j.neuroimage.2016.08.016

74. Vizioli, L., Moeller, S., Dowdle, L., Akçakaya, M., De Martino, F., Yacoub, E., Uğurbil, K., 2021. Lowering the thermal noise barrier in functional brain mapping with magnetic resonance imaging. Nat. Commun. 12, 5181. https://doi.org/10.1038/s41467-021-25431-8

75. Vu, V.Q., Ravikumar, P., Naselaris, T., Kay, K.N., Gallant, J.L., Yu, B., 2011. Encoding and Decoding V1 Fmri Responses to Natural Images with Sparse Nonparametric Models. Ann Appl Stat 1159–1182.

76. Wald, L.L., Polimeni, J.R., 2017. Impacting the effect of fMRI noise through hardware and acquisition choices - Implications for controlling false positive rates. Neuroimage 154, 15–22. https://doi.org/10.1016/j.neuroimage.2016.12.057

77. Warren, S.G., Yacoub, E., Ghose, G.M., 2014. Featural and temporal attention selectively enhance task-appropriate representations in human primary visual cortex. Nat. Commun. 5, 1–12. https://doi.org/10.1038/ncomms6643

78. Weldon, K.B., Olman, C.A., 2021. Forging a path to mesoscopic imaging success with ultra-high field functional magnetic resonance imaging. Philos. Trans. R. Soc. B Biol. Sci. 376, 20200040. https://doi.org/10.1098/rstb.2020.0040

79. Yacoub, E., Duong, T.Q., Moortele, P.-F.V.D., Lindquist, M., Adriany, G., Kim, S.-G., Uğurbil, K., Hu, X., 2003. Spin-echo fMRI in humans using high spatial resolutions and high magnetic fields. Magn. Reson. Med. 49, 655–664. https://doi.org/10.1002/mrm.10433

80. Yacoub, E., Harel, N., Uğurbil, K., 2008. High-field fMRI unveils orientation columns in humans. Proc. Natl. Acad. Sci. 105, 10607–10612. https://doi.org/10.1073/pnas.0804110105

81. Zaretskaya, N., 2021. Zooming-in on higher-level vision: High-resolution fMRI for understanding visual perception and awareness. Prog. Neurobiol. 207, 101998. https://doi.org/10.1016/j.pneurobio.2021.101998

82. Zhao, F., Wang, P., Hendrich, K., Ugurbil, K., Kim, S.G., 2006. Cortical layer-dependent BOLD and CBV responses measured by spin-echo and gradient-echo fMRI: insights into hemodynamic regulation. NeuroImage 1149–1160.

